# Predictive profiling of SARS-CoV-2 variants by deep mutational learning

**DOI:** 10.1101/2021.12.07.471580

**Authors:** Joseph M. Taft, Cédric R. Weber, Beichen Gao, Roy A. Ehling, Jiami Han, Lester Frei, Sean W. Metcalfe, Alexander Yermanos, William Kelton, Sai T. Reddy

## Abstract

The continual evolution of the severe acute respiratory syndrome coronavirus-2 (SARS-CoV-2) and the emergence of variants that show resistance to vaccines and neutralizing antibodies (*1–4*) threaten to prolong the coronavirus disease 2019 (COVID-19) pandemic (*5*). Selection and emergence of SARS-CoV-2 variants are driven in part by mutations within the viral spike protein and in particular the ACE2 receptor-binding domain (RBD), a primary target site for neutralizing antibodies. Here, we develop deep mutational learning (DML), a machine learning-guided protein engineering technology, which is used to interrogate a massive sequence space of combinatorial mutations, representing billions of RBD variants, by accurately predicting their impact on ACE2 binding and antibody escape. A highly diverse landscape of possible SARS-CoV-2 variants is identified that could emerge from a multitude of evolutionary trajectories. DML may be used for predictive profiling on current and prospective variants, including highly mutated variants such as omicron (B.1.1.529), thus supporting decision making for public heath as well as guiding the development of therapeutic antibody treatments and vaccines for COVID-19.

## Introduction

As of late 2021, variants of SARS-CoV-2 associated with higher transmissibility and/or immune evasion (antibody escape) have almost entirely supplanted the original founder strain (Wu-Hu-1) (*6*). Emerging variants often possess at least one mutation in the RBD (*1, 7–9*), which can directly influence binding to ACE2 (*10, 11*). For example, alpha (B.1.1.7), beta and gamma variants all possess the N501Y mutation, which is associated with higher affinity binding to ACE2 (*12*), suggesting this may represent a possible selective pressure for variant emergence.

Neutralizing antibodies, including monoclonal antibody therapeutics and those induced by vaccination (with the original Wu-Hu-1 spike protein), often display reduced binding and neutralization to variants. Detailed molecular analysis has revealed that many neutralizing antibodies to SARS-CoV-2 share sequence and structural features (*13–17*), which has led to their categorization into four common classes defined by groups of targeted RBD epitopes (*16, 18*). For example, class 1 antibodies include the clinically approved REGN10933 (casirivimab) (*19, 20*) and LY-CoV16 (etesevimab) (*21*). Circulating variants with mutations in position K417 [e.g., beta, gamma and delta plus (B.1.617.2 + K417N)] as well as the mink-selected Y453F mutation (Cluster 5) display decreased neutralization by these class 1 antibodies (*1, 2, 22*). Class 2 neutralizing antibodies including the clinically used LY-CoV555 (bamlanivimab) also strongly inhibit ACE2 binding, however, variants such as beta, gamma, eta (B.1.525), kappa (B.1617.1) and iota (B1.526) all possess the RBD mutation E484K/Q that can lead to a substantial loss of binding and neutralization. Class 3 antibodies, including the clinically approved REGN10987 (imdevimab) and S309 (sotrovimab) (*23*), bind partially conserved epitopes and have so far retained binding to most circulating variants, with the exception of the N439K mutation, which reduces binding of REGN10987 by 28-fold (*24*). Class 4 antibodies such as CR3022 target a highly conserved epitope among sarbecoviruses (*25–27*), and are therefore largely resistant to escape variants, but generally lack neutralizing potency since they do not directly inhibit ACE2 binding. This clustering of human neutralizing antibodies into discrete epitope-binding classes further suggests that the evolution of SARS-CoV-2 may converge via mutations that provide escape to these common antibody classes, which could result in variants with immune evasion.

Bloom and colleagues have performed yeast surface display and deep mutational scanning (DMS) (*28*) on the entire 201 amino acid RBD of SARS-CoV-2 in order to determine the impact of single-position substitutions on binding to ACE2 and monoclonal or serum antibodies (*22, 29–32*). While DMS has been very effective at single mutation profiling of the RBD, several widely circulating variants (e.g., beta, gamma and delta) as well as newly emerging variants [e.g., mu (B.1621) and lambda (C.37)] possess multiple mutations in the RBD, which are associated with enhanced ACE2 binding and/or multi-class antibody escape (*18*). Critically, the recent emergence of the omicron variant possessing 15 RBD mutations represents a substantial risk for immune evasion, thus underscoring the urgent need to determine the impact of combinatorial mutations. However, combinatorial sequence space grows exponentially as the number of mutations and amino acid diversity increases, rapidly outpacing the capabilities of experimental screening techniques. For instance, when focusing only on a subset of twenty RBD residues directly involved in ACE2 binding (*33*), theoretical sequence space (20^20^ = 1 x 10^26^) far exceeds what can be screened by yeast display libraries (~10^9^). Here, we establish deep mutational learning (DML), which integrates experimental yeast display screening of RBD mutagenesis libraries with deep sequencing and machine learning (**Fig. 1**). DML provides comprehensive interrogation of combinatorial RBD mutations and their impact on ACE2 binding and antibody escape, thus enabling predictive profiling of SARS-CoV-2 variants.

**Figure 1.**
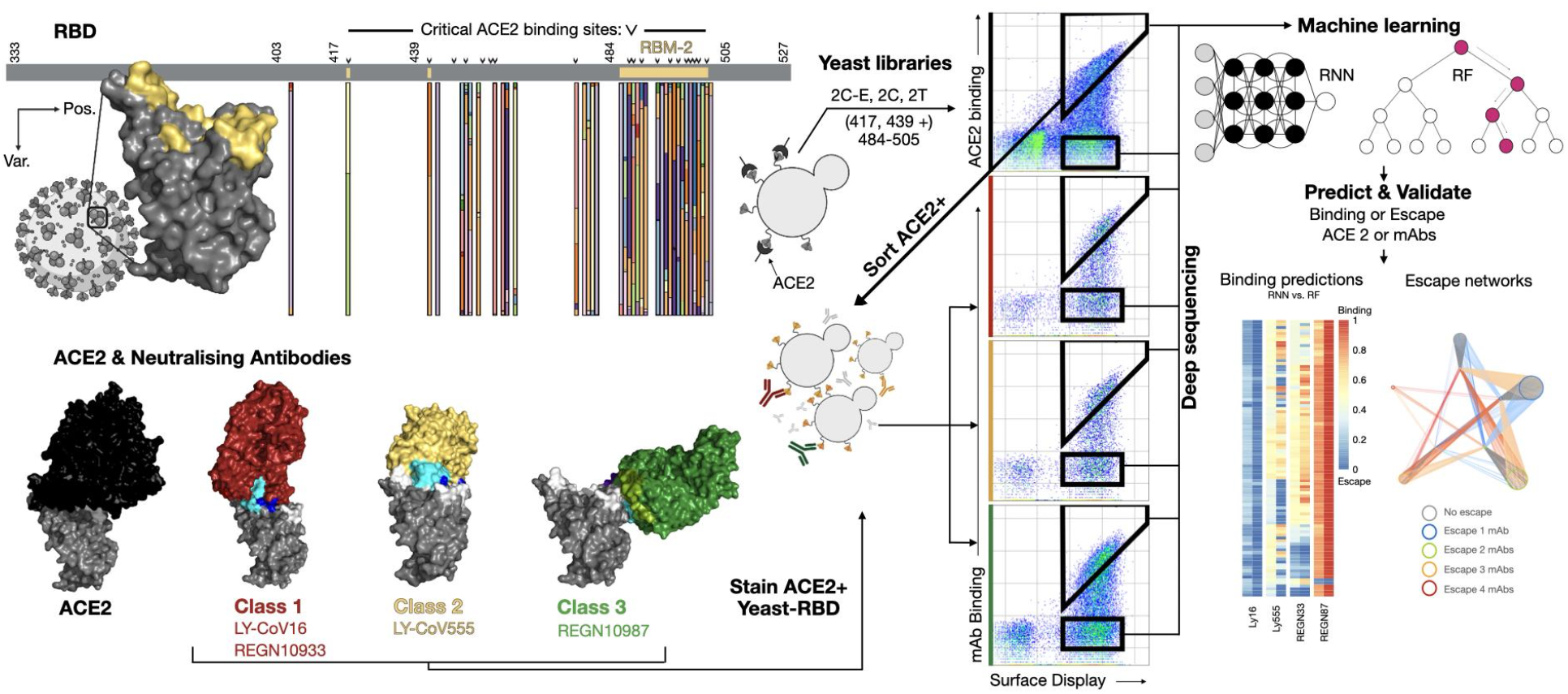
Overview of deep mutational learning of the RBD for prediction of ACE2 binding and antibody escape. The RBD or the SARS-CoV-2 spike protein is expressed on the surface of yeast, mutagenesis libraries are designed on the receptor-binding motif (RBM-2) of the RBD, which are the sites of interaction with ACE2 and neutralizing antibodies (e.g., therapeutic antibody drugs). RBD libraries are screened by FACS for binding to ACE2 and neutralizing antibodies, both binding and non-binding (escape) populations are isolated and subjected to deep sequencing. Machine learning models are trained to predict binding status to ACE2 or antibodies based on RBD sequence. Machine learning models are then used to predict ACE2 binding and antibody escape on current and prospective variants and lineages.

### Design and screening of RBD mutagenesis libraries

SARS-CoV-2 RBD mutagenesis libraries were targeted to a core region of the receptor-binding motif (RBM-2, positions 484-505), a subregion of the RBD that directly interfaces with ACE2 and where mutations are commonly observed in viral genome sequencing data [available on GISAID (www.gsaid.org)]. To generate training datasets covering a high mutational sequence space, a combinatorial mutagenesis scheme was designed based on DMS data for ACE2 binding, previously published by Starr et al. (*29*). Single mutation fitness values were empirically thresholded and converted to amino acid frequencies, with mutations below the ACE2 binding fitness threshold excluded. For each position, degenerate codons approximating the desired amino acid distribution were selected by minimizing mean-squared error (*34*) (some positions remained fixed due to their inability to tolerate mutations and retain ACE2 binding), resulting in a library with a theoretical amino acid diversity of 1.50 x 10^10^ (library 2C) (**Fig. 2a**). An extended version of this library was also designed, with fully degenerate codons (NNK) at positions 417 and 439, which are mutated in a number of circulating variants and associated with antibody escape (*35, 36*), resulting in a theoretical amino acid diversity of 5.99 x 10^12^ (library 2CE). To generate training datasets covering a lower mutational sequence space, we constructed a tiling mutagenesis library, whereby fully degenerate codons (NNK) were tiled across three of the positions in RBM-2, resulting in a theoretical amino acid diversity of 1.53 x 10^6^ (library 2T) (**Fig. 2b**).

**Figure 2.**
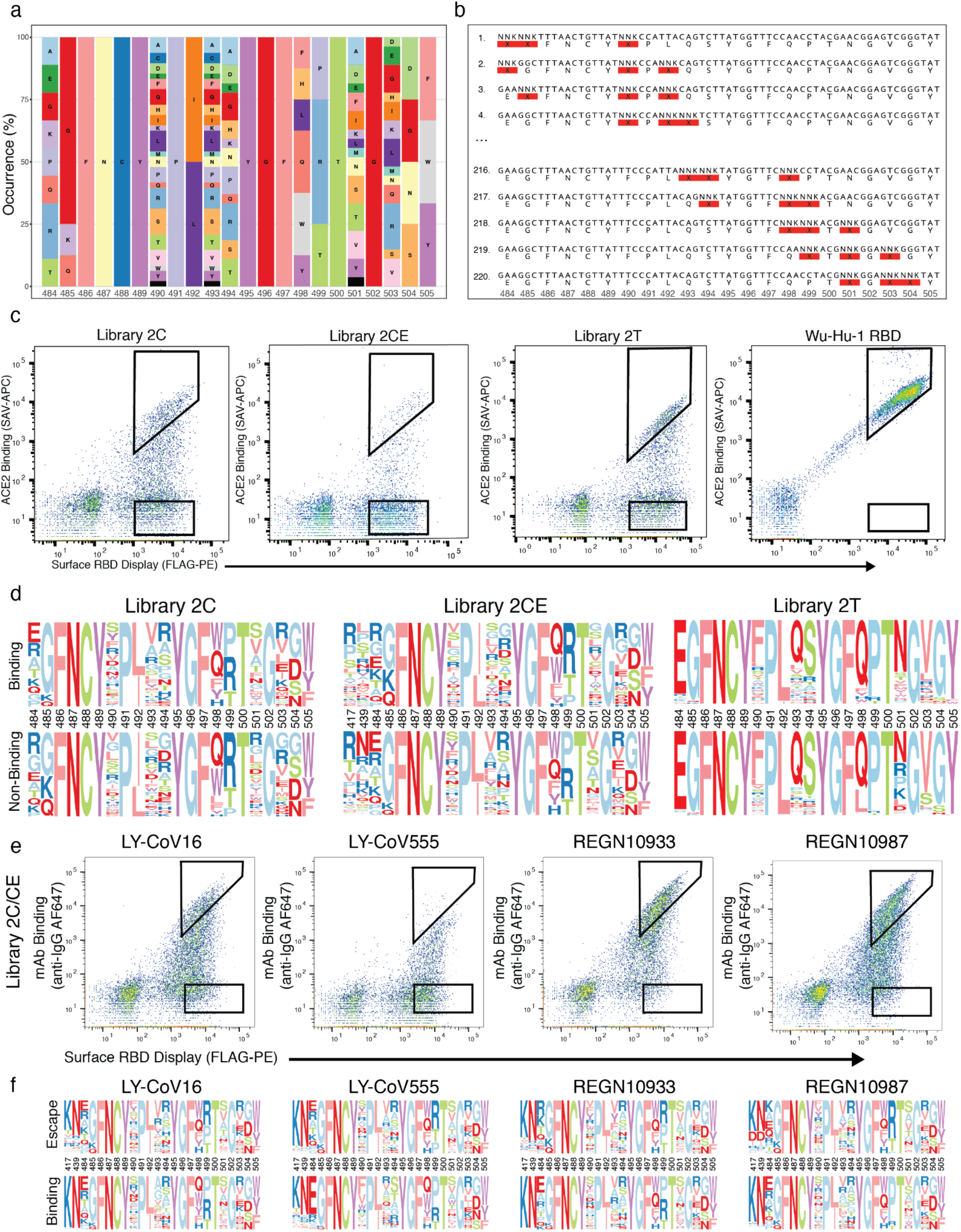
Design of RBD mutagenesis libraries and screening by yeast surface display and deep sequencing. **a,** Shown is the amino acid usage in positions RBM-2 (484-505 of the RBD), which was based on DMS data for ACE2 binding (*29*) and used to design the combinatorial library 2C. **b,** Shown are representative examples of degenerate codons tiled across RBM-2, which are pooled together to comprise library 2T. **c,** Flow cytometry dot plots depict yeast display screening of RBD libraries (2C, 2CE and 2T) and control RBD (Wu-Hu-1); gating schemes correspond to selection of ACE2-binding and non-binding variants. **d,** Amino acid logo plots of the RBD are based on deep sequencing data from ACE2-binding and non-binding selections. **e,** Flow cytometry dot plots depict yeast display screening of pooled RBD libraries (2C and 2CE) that were pre-selected for ACE2 binding; gating schemes correspond to selection of variants for binding and escape (non-binding) to four therapeutic monoclonal antibodies (mAbs). **f,** Amino acid logo plots of the RBD are based on deep sequencing data from antibody-binding and escape selections.

Synthetic oligonucleotides encoding the different libraries and spanning the region of interest were amplified by PCR to produce double-stranded DNA with homology to the full RBD sequence. Co-transformation of yeast (*S. cerevisiae* EBY100) with library-encoding DNA and linearized plasmid yielded more than 2 x 10^7^ transformants for each library. RBD variants, displayed on the yeast surface as a C-terminal fusion to Aga2 (*37*), were isolated by fluorescence-activated cell sorting (FACS) based on binding to soluble human ACE2 receptor (Wu-Hu-1 RBD used as a guide for gating). RBD variants which showed a complete loss of binding to ACE2 were also isolated (**Fig. 2c, Supplementary Table 1**). Importantly, this did not include variants with only partially reduced binding since such an intermediate population could not be assigned as binding or non-binding with sufficient confidence necessary for training supervised machine learning models. Targeted deep sequencing (Illumina) of the RBD gene was performed on all the sorted libraries; protein sequence logos revealed highly similar patterns of amino acid usage between the ACE2-binding and non-binding fractions (**Fig. 2d, Supplementary Table 2**).

Next, using exclusively the ACE2-binding populations, FACS was performed to isolate variants that maintained binding or showed a complete loss of binding (escape) to four clinically used antibodies (LY-CoV16, LY-CoV555, REGN10933 and REGN10987). Wu-Hu-1 RBD was again used as a guide for gating antibody binding or escape (**Fig. 2e, Supplementary Fig. 1**). The proportion of binding and escape (non-binding) for each antibody and library was highly variable, with REGN10933 having the lowest fraction of escape variants and LY-CoV555 having the highest (**Supplementary Table 1**). Deep sequencing was once again performed on antibody binding and escape fractions of all the sorted RBD libraries, and similar to ACE2, protein sequence logos of the two fractions looked highly similar (**Fig. 2f**).

### Machine learning models accurately predict ACE2 binding and antibody escape

Deep sequencing data from ACE2 selections underwent pre-processing, quality filtering and balancing steps to create the final training sets (**Methods, Supplementary Tables 3, 4**). Following nucleotide to protein translation, amino acid sequences were converted to an input matrix by one-hot encoding (**Fig. 3a**). Supervised machine learning models were trained for classification of ACE2 binding, which is defined as the probability (*P*) that any given RBD sequence binds to ACE2 (higher *P* correlates with binding). For initial benchmarking, a range of different baseline models (default parameters) were evaluated for their classification performance across several metrics (accuracy, F1, precision, recall). Machine learning models tested included K-nearest neighbor, logistic regression, naive Bayes, support vector machines, and random forests (RF); long-short term memory recurrent neural networks (RNN) were also trained, which are a class of deep learning models that have the ability to learn long-range dependencies in sequential data (*38–41*).

**Figure 3.**
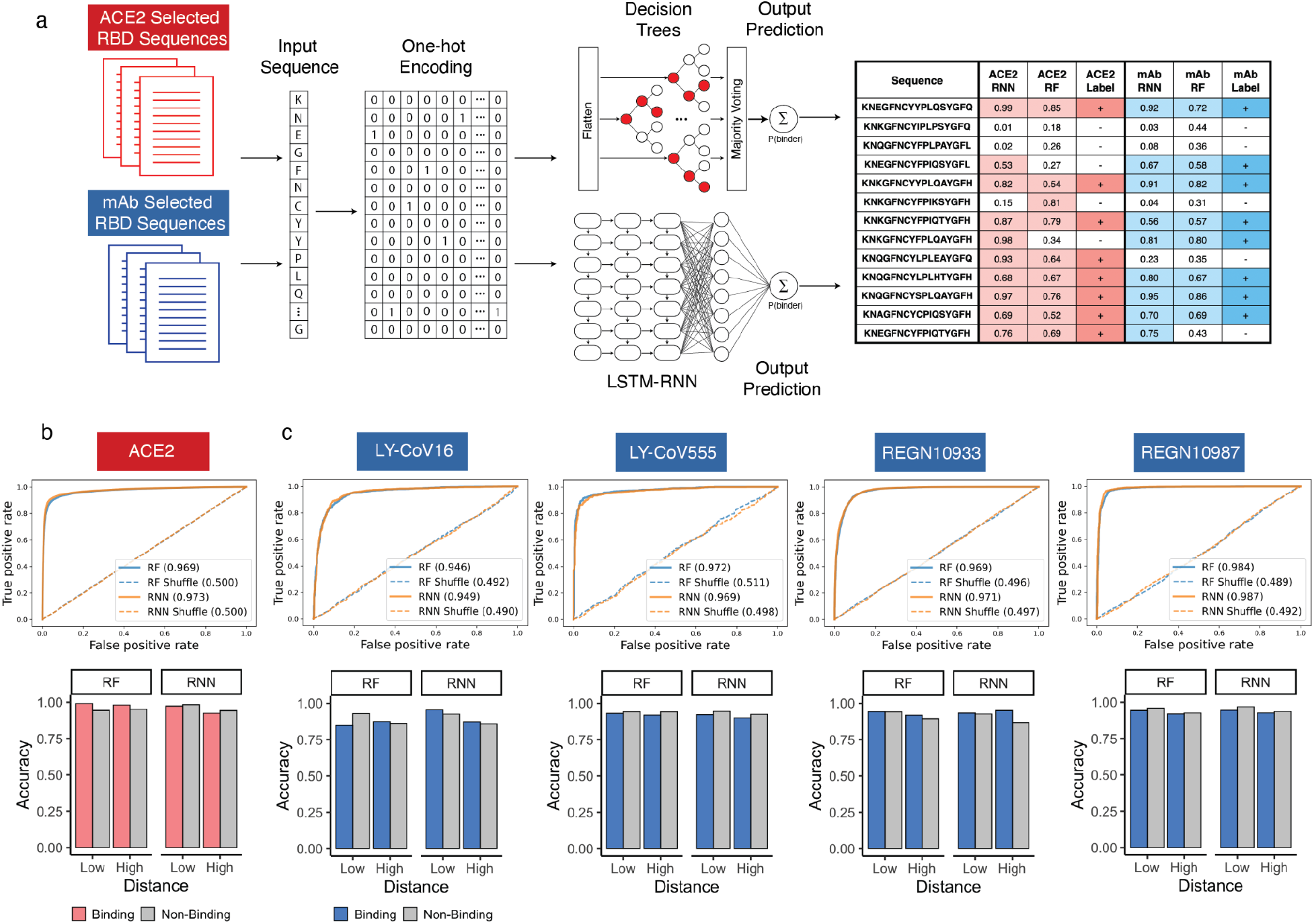
Training and testing of machine and deep learning models for prediction of ACE2 binding and antibody escape based on RBD sequence. **a,** Deep sequencing data from ACE2 and monoclonal antibody (mAb) selections is encoded by one-hot encoding and used to train supervised machine learning (e.g., Random Forest, RF) and deep learning models (e.g., recurrent neural network, RNN). Models perform classification by predicting a probability (*P*) of ACE2 binding or non-binding and mAb binding or escape (non-binding) based on the RBD sequence. **b** and **c,** Performance of fully trained and optimized RF and RNN models for prediction of ACE2 binding and antibody escape on test data is shown with ROC curves and accuracy graphs. Low and high distance sequences are defined as those ≤ ED_5_ and ≥ ED_6_ from Wu-Hu-1 RBD, respectively.

While all baseline models performed effectively (e.g., accuracy scores between 0.87 - 0.94), RF and RNN were selected for further optimization and application since they showed relatively higher performance metrics and could be trained faster **(Supplementary Fig. 2)**. After hyperparameter optimization, the classification performance of both RF and RNN models was improved further, resulting in an area under ROC curve (AUC) values of 0.969 and 0.973, respectively (**Fig. 3b, Supplementary Table 5**).

SARS-CoV-2 is evolving across a range of mutational trajectories, including variants such as omicron that rapidly accumulate mutations, as well variants that develop into lineages with subvariants (e.g., C.1, C.1.1, C.1.2) (*42, 43*). Determining the performance of machine learning models across various mutational edit distances [Edit Distance (ED) from the reference Wu-Hu-1 RBD sequence] is therefore an important criterion. We thus examined model performance on test data that was divided into low mutational distances (≤ ED_5_), which corresponds to variants such as beta and gamma, and high mutational distances (≥ ED_6_), which corresponds to variants such as omicron. For both low and high distance variants, RF and RNN models showed high accuracy scores (>94% and >92% for low and high distances respectively) (**Fig. 3b, Supplementary Fig. 3a, Supplementary Tables 6, 7**).

Similar to the ACE2 selections, deep sequencing data from antibody selections was pre-processed, quality filtered, balanced and encoded as before. Supervised machine learning models (RF and RNN) were trained to classify antibody escape, which is defined as the probability that a given RBD sequence escapes a defined antibody (lower *P* correlates with escape). Following hyperparameter optimization, individual models trained for each of the four therapeutic antibodies displayed high performance metrics (**Fig. 3c**). Similar to ACE2 models, antibody escape models also showed high performance metrics across both low and high distance variants (**Supplementary Tables 6, 7**). Closer examination revealed that it was crucial to combine training data from low distance libraries (2T) and high distance libraries (2C, 2CE) to achieve better performance for ACE2 and antibody escape models (**Supplementary Fig. 3b**).

### Predictive profiling on synthetic lineage variants

Having established that ACE2 binding and antibody escape machine learning models can make highly accurate predictions on test data, we next evaluated their classification performance on defined variants, followed by experimental validation and structural modeling (**Fig. 4a**). Synthetic lineages were generated *in silico* to simulate plausible evolutionary paths, where variants without predicted ACE2-binding intermediates at each mutational step were excluded. The lineages were designed to include variants at ED_3_, ED_5_ and ED_7_ from the original Wu-Hu-1 RBD sequence (nucleotide and amino acid). Additionally, the sequences were chosen to form lineages containing mutations observed in circulating variants (e.g., alpha: N501Y, beta/gamma: E484K and N501Y, kappa: E484Q and N501Y). ACE2 binding was predicted based on a consensus model, whereby a given RBD sequence is predicted to bind ACE2 when both RF and RNN models yield *P* > 0.5, else they are predicted to be non-binders. The 46 synthetic lineage variants were chosen to contain diversity in ACE2 binding prediction (36 predicted binders, 10 predicted non-binders) (**Fig. 4b**). Additionally, predictions for escape from each of the four therapeutic antibodies were made for the synthetic variants using a similar consensus model approach (RBD sequence escapes an antibody when both RF and RNN outputs are *P* < 0.5) (**Supplementary Fig. 4a–d**). After having made all machine learning predictions, each synthetic RBD variant was individually expressed on the surface of yeast cells and assessed for ACE2 binding and antibody escape. The consensus model correctly predicted ACE2 binding for 91.67% (33/36) of the synthetic variants, with an accuracy of non-binding prediction of 100% (10/10), resulting in an overall prediction accuracy of 93.48% (43/46) (**Fig. 4b, Supplementary Fig. 5**). For the 33 correctly predicted ACE2-binding variants, the combined accuracy of antibody escape predictions across all four therapeutic antibodies was 93.94% (124/132) (LY-CoV16: 31/33, LY-CoV555: 30/33, REGN10933: 31/33, REGN10987: 32/33) (**Fig. 4c, Supplementary Fig. 4a–d**). Additionally, we identified three variants that were only ED3 (nucleotide and amino acid) from the Wu-Hu-1 RBD and in which consensus models predicted ACE2 binding and escape from all four therapeutic antibodies. One of these variants possessed mutations in positions 493, 498 and 501, which are all mutated in the omicron variant. Subsequent yeast display experiments confirmed these machine learning predictions of antibody escape to all four therapeutic antibodies, including escape from the often mutation-resistant REGN10987 (**Supplementary Fig. 6**). Structural modeling by AlphaFold2 (*44*) was performed on eight synthetic RBD variants (all variants were accurately classified and experimentally validated for ACE2 binding or non-binding) (**Fig. 4d**). The structural predictions showed that several ACE2 non-binding variants did not differ substantially from the original Wu-Hu-1 RBD. In contrast, the ACE2-binding variants showed a wide diversity of possible structural conformations.

**Figure 4.**
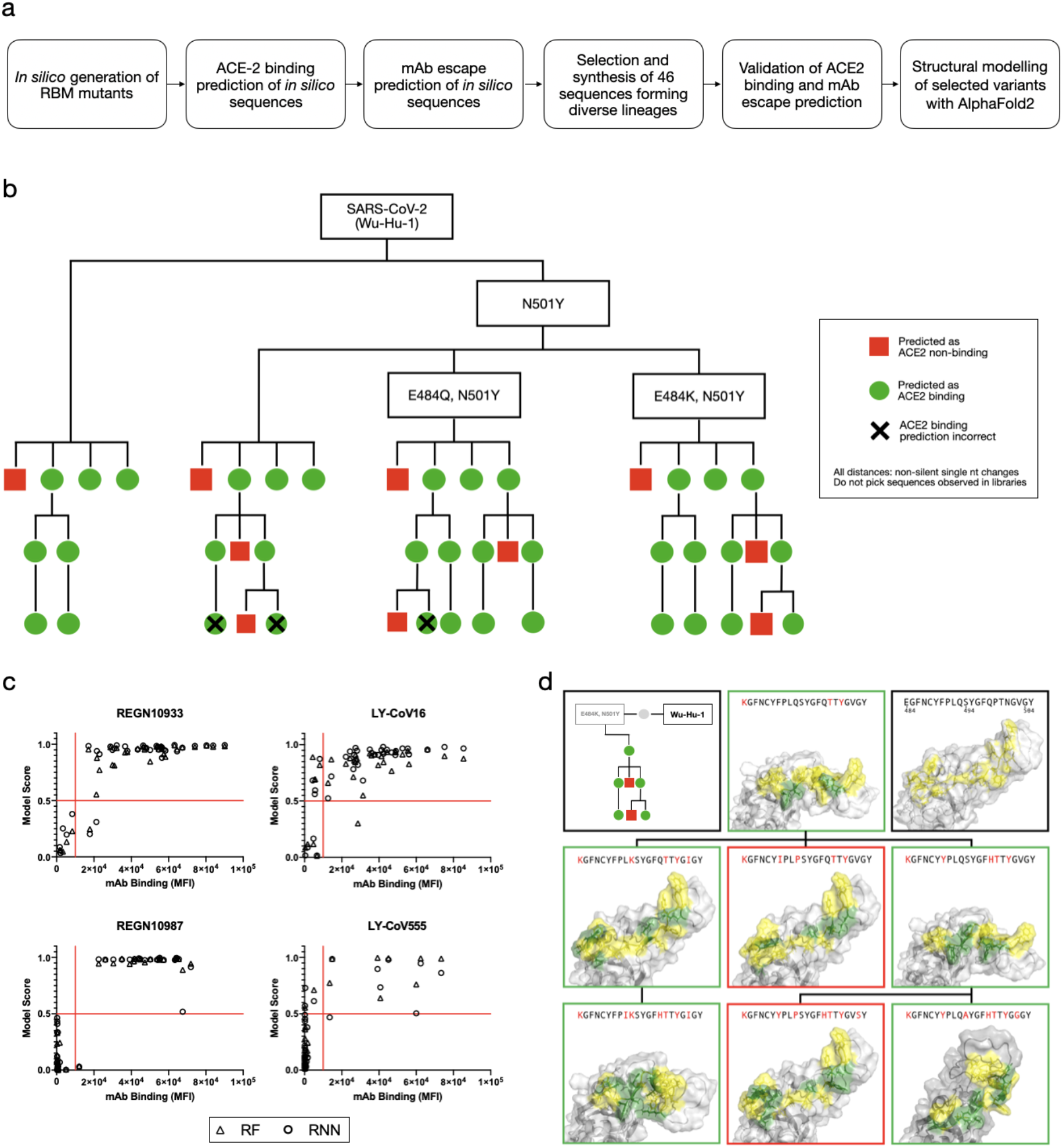
Prediction and experimental validation of synthetic lineages of RBD variants. **a,** Workflow to select and test synthetic variants at chosen edit distances (ED_3_, ED_5_, and ED_7_) from Wu-Hu-1 RBD. **b,** Lineage plot of synthetic variants depicts machine learning predictions and experimental validation (**Supplementary Fig. 5**) for ACE2 binding and non-binding. **c,** Dot plots of synthetic variants correspond to machine learning model (RF and RNN) predictions and experimental validation for antibody binding or escape. **d,** Structural modeling by AlphaFold2 shows predicted structures of RBD variants that are ACE2 binding (green boxes) or non-binding (red boxes); control is Wu-Hu-1 RBD (black box).

### Predictive profiling of current and prospective variants

In addition to the selected synthetic lineages, we also performed machine learning to predict ACE2 binding and antibody escape on a panel of 12 naturally-occurring variants of SARS-CoV-2 (**Supplementary Table 8**). We determined the accuracy of machine learning predictions on antibody escape by using, as a reference, the Stanford SARS-CoV-2 Susceptibility Database (*24*). Applying the same consensus model approach (RF and RNN) and thresholds as before, the prediction accuracy for ACE2 binding was 100% (12/12) and the prediction accuracy for escape across all four therapeutic antibodies was 85.42% (41/48) (antibody escape is defined here as a reported 30-fold reduced neutralization in the Stanford database). Strikingly, when applying a more stringent threshold for antibody escape prediction that requires both RF and RNN models to have high certainty in their prediction (both models *P* < 0.25 for escape and *P* > 0.75 for binding), 100% (30/30) of the machine learning predictions matched the results reported in the Stanford database.

Next, we used machine learning models to predict antibody escape on prospective ACE2-binding lineages at low mutational distances (ED_1_ and ED_2_) from the Wu-Hu-1, alpha, beta, kappa, gamma and B.1.1523 RBD sequences. (**Fig. 5, Supplementary Figs. 7–9, Supplementary Table 9**). Using a stringent threshold for antibody escape, we identified distinct patterns based on the starting variant. For example, REGN10933 and REGN10987 were largely resilient to escape from ED_1_ lineages of Wu-Hu-1, alpha and kappa (**Fig. 5a–i and Supplementary Fig. 7a–i**). While ED_1_ lineages of beta and gamma almost entirely escape both LY-CoV555 and LY-CoV16. A large fraction of ED2 lineages from all variants escaped REGN10933, LY-CoV555 and LY-CoV16, revealing an increasing likelihood of escape with an increasing number of mutations. Notably, a small fraction (0.17%) of beta ED2 lineages are predicted to escape all four therapeutic antibodies, whereby several of these variants possessed mutations in positions 417, 484, 493 and 501, which are all mutated in the omicron variant (**Fig. 5f**). For further visualization, we constructed deep escape networks (**Fig. 5c, f, i** and **Supplementary Fig. 7c, f, i**), depicting the vulnerability of the four therapeutic antibodies to low distance mutations (ED_1_ and ED_2_). Specifically, deep escape networks illustrate the increase in sequence space per mutation while also pointing out the presence of mutations that vastly increase escape from multiple antibodies. For example, there are variants at ED_1_ from Wu-Hu-1 that are predicted to not escape any of the four antibodies, however, just one additional mutation (ED_2_) can result in variants predicted to escape up to three antibodies.

**Figure 5.**
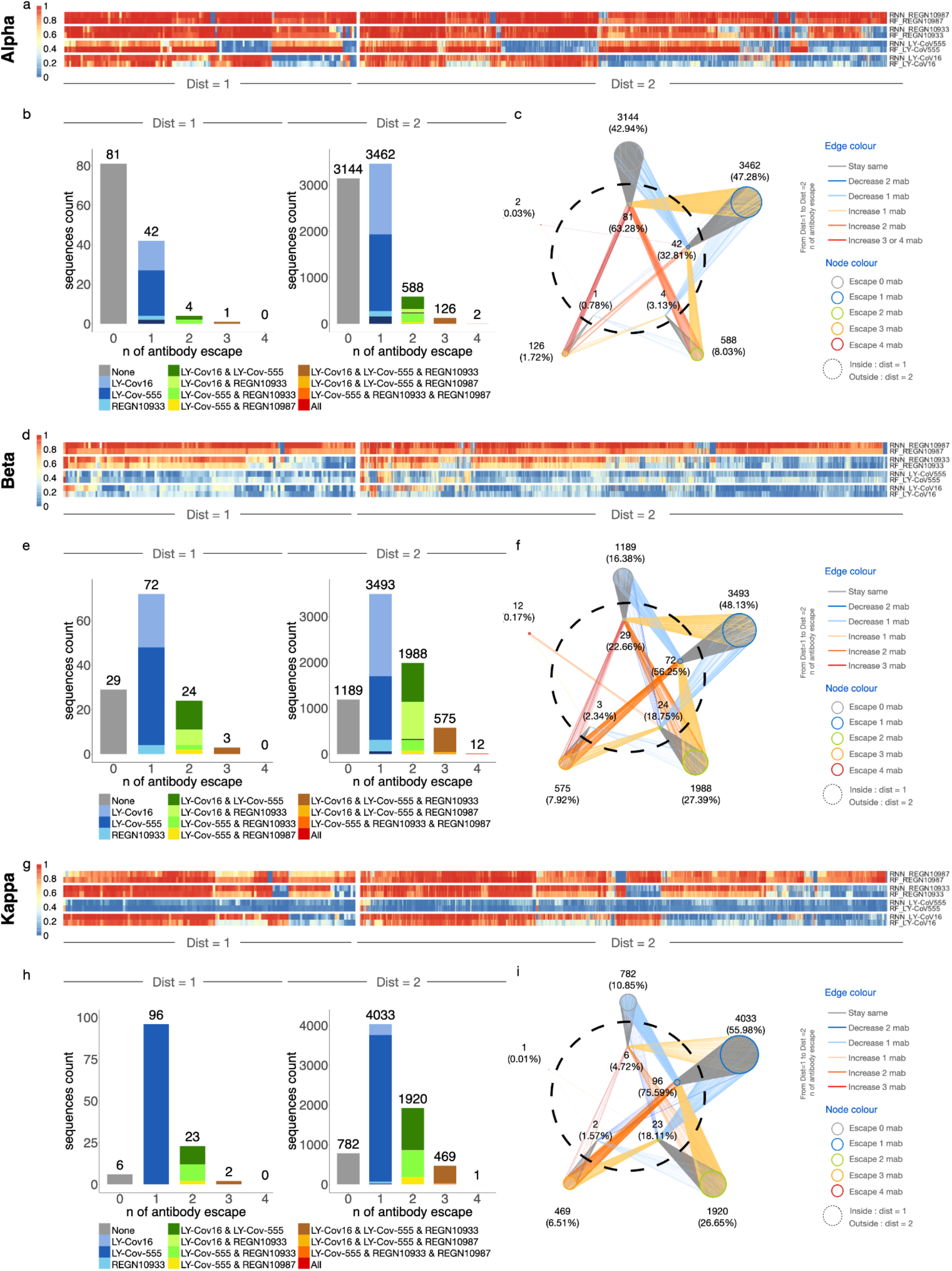
Predictive profiling of selected RBD variants for antibody escape across low mutational distances. **a, d, g,** Heatmap depicts monoclonal antibody (mAb) binding as assessed by RF and RNN models of ED_1_ and ED_2_ variants of alpha, beta and kappa. **b, e, h,** The number of sequences escaping a combination of *n* (number) mAbs for ED_1_ and ED_2_ (agreement between models, threshold >0.5). **c, f, i,** Deep escape networks display possible evolutionary paths between variants and their escape from mAbs.

Finally, DML enables rapid *in silico* evaluation of new variants that appear on genomic databases (GISAID). For example, we performed a similar analysis on the recently reported B.1.1.523 variant possessing RBD mutations E484K and S494P (*45*), which revealed complete escape from LY-CoV555 and ED_1_ lineages, as well as substantial escape for other antibodies in ED_2_ lineages, including three variants that escaped all four therapeutic antibodies (**Supplementary Fig. 7g, h, i**).

## Discussion

For a multitude of reasons, eradication of SARS-CoV-2 appears improbable. Instead, an endemic future likely awaits (*46, 47*). An endemic and continually evolving SARS-CoV-2 poses a perpetual risk for the emergence of new variants that escape from vaccine-or infection-induced antibodies. In this study, we develop DML, a machine learning-guided protein engineering method for determining the impact of mutations in the SARS-CoV-2 RBD on ACE2 binding and antibody escape. In DML, machine learning models trained on thousands of classified RBD variants obtained from library screening make highly accurate predictions across a sequence space of billions of RBD variants, several orders of magnitude larger than what is possible from experimental screening alone. While we focused in this study on a subregion of the RBD associated with variants such as alpha, beta and gamma, this method can be expanded to incorporate mutations in additional regions across the RBD, which are associated with other variants and their corresponding lineages (e.g., omicron, delta, delta plus) (*48*).

DML could be used as a de facto monitoring system by rapidly and efficiently making predictions on genomic surveillance data of new variants, without the immediate need for experimental assays. This is especially crucial given the emergence of highly mutated variants such as omicron, where crucial public health decisions have to urgently be made, often before experimental assays of antibody escape or vaccine resistance can be performed.

By providing accurate predictions of antibody escape across a large mutational landscape, DML may enable researchers to select candidate antibody therapeutics and cocktails with the broadest efficacy against the spectrum of possible variants, some of which may occur simultaneously and may be highly mutated such as omicron. Assessing the efficacy of candidate antibodies against future variants puts therapeutic development on a proactive rather than reactive footing, potentially avoiding cases like LY-CoV555, which has lost efficacy to most variants of concern (*22, 24*), resulting in the revocation of its clinical authorization as a monotherapy. Furthermore, such an approach could be used to guide the development of antibodies and cocktails that maximize breadth and potency (*49, 50*) to both current and prospective variants. Finally, evidence exists that the receptor-binding domains of other endemic coronaviruses may be undergoing adaptive evolution to escape from human antibody responses (*51, 52*). Consequently, the application of DML to predict SARS-CoV-2 escape from polyclonal antibodies present in serum of vaccinated or convalescent individuals, combined with phylogenetic models of viral evolution (*53*), may enable the prospective identification of future variants with the highest likelihood of emergence and thus support vaccine development for COVID-19.

## Supporting information

STable 3

STable 4

STable 6

STable 7

STable 8

STable 9

## Data and materials availability

The main data supporting the results in this study are available within the paper and its Supplementary Information.

## Acknowledgments

We thank the ETH Zurich D-BSSE Single Cell Unit and the Genomics Facility Basel for support.

## Author contributions

Conceptualization: JMT, CRW, BG, RAE and STR. Experimental methodology: JMT, RAE and LF. Computational methodology: JMT, CRW, BG, SWM, JH, AY. Supervision: JMT, CRW and STR. Writing - original draft: JMT, CRW, BG, RAE and STR. Writing - review & editing: all authors.

## MATERIALS & METHODS

### Rational design of SARS-CoV-2 RBD mutagenesis libraries

#### Combinatorial library design

The design of the combinatorial library 2C consisted of mutating residues within the RBM-2 region (positions 484-505 of the RBD) and was based on previously described results from deep mutational scanning (DMS) experiments (*1*). DMS enrichment ratios described by Starr et al. were thresholded to exclude mutations with decreased ACE2-binding fitness and then converted to amino acid frequencies as described previously (*2*). For each position, degenerate codons approximating the amino acid frequency distribution and diversity were selected, resulting in a library with a theoretical diversity of 1.50 x 10^10^ amino acid sequences. Library 2CE consisted of the same combinatorial design in positions 484-505 but with additional fully degenerate codons (NNK) in positions 417 and 439, resulting in a theoretical amino acid diversity of 5.95 x 10^12^.

#### Tiling library design

The tiling library 2T was designed by incorporating three positions with full degenerate codons (NNK) within the RBM-2 (positions 484-505) of the RBD. The degenerate codons are tiled across such that the total sequences of a tiling library, i.e., the number of variants of up to a maximum edit distance (ED) *k* away from the wild-type sequence is determined by the length of sequence (*n*, here = 14 non-fixed positions in RBM-2), the number of NNKs (or max ED) introduced (*k*, here = 3) and the size of the alphabet (*a*, here = 20):

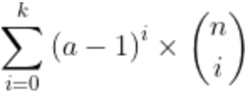

Similarly, the number of sequences for a given ED *k* is given by:

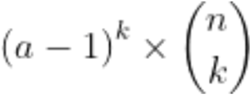

The resulting total diversity of the library 2T is 1,533,035 sequences.

### Cloning and expression of RBD mutagenesis libraries for yeast surface display

For libraries 2C and 2CE, synthetic single-stranded oligonucleotides (ssODNs) (Integrated DNA Technologies ultramers or oPools) were designed with degenerate codons spanning the region of interest and encoding the desired library diversity, with 30 bp overhangs on each end that were homologous to the yeast display plasmid pYD1. For library 2T, pools of ssODN were designed, where each member of the pool contains one combination of the three ‘NNK’ codons; in this case, consisting of 120 unique ssODNs. The ssODNs were amplified by PCR to produce doublestranded DNA. The plasmid pYD1 was modified such that the entire C-terminal fusion to Aga2 was replaced with a cassette encoding the RBD (Wu-Hu-1 sequence), expression tags and stop codon (HA Tag-RBD-FLAG-Stop). The RBM-2 residues 484-505 were replaced with an EcoRI recognition site, allowing production of a linearized vector with homology to mutagenesis ssODNs and with no parental background. Insert and EcoRI-linearized plasmids were concentrated and purified by silica spin columns (Zymo D4013) followed by drop dialysis for 1 hour in nuclease-free H2O (Millipore VSWP02500). The libraries were cloned and expressed in yeast by *in vivo* homologous recombination, as previously described (*3, 4*), using 1 μg each of plasmid and insert DNA per 300 μl of electrocompetent EBY100 cells in a 2 mm electroporation cuvette.

### Screening RBD libraries for ACE2-binding and non-binding

Surface expression of SARS-Cov2 RBD was induced by growth in SG-UT medium at 23°C for 16-40 hours, as previously described (*3*). Approximately 10^8^ library cells were washed once with 1 mL wash buffer (Dulbecco’s PBS+ 0.5% BSA + 0.1% Tween20 + 2 mM EDTA) by centrifugation at 8000 x g for 30 s. Washed cells were stained with 50 nM biotinylated human ACE2 (Acros AC2-H82E6) for 30 minutes at 4 °C, followed by an additional wash. Cells were then stained with 2.5 ng/μlstreptavidin-AlexaFluor 647 (Biolegend 405237) and 1 ng/μl anti-FLAG-PE (Biolegend 637310) for 30 minutes at 4 °C. Cells were subsequently pelleted by centrifugation at 8000 x g for 30s and kept on ice until sorting. Binding (ACE2+/FLAG+) and non-binding (ACE2-/FLAG+) cells were sorted by FACS (BD FACSAria Fusion or Sony MA800 cytometer) (**Fig. 2**). Collected cells were cultured in SD-UT medium for one to two days at 30 °C. Induction and sorting was repeated until the desired populations were pure.

### Screening RBD libraries for binding and escape to monoclonal antibodies

RBD libraries pre-sorted for ACE2-binding were cultured and induced, as described above. Induced cells were washed once with DPBS wash buffer, followed by incubation with 100 nM monoclonal antibody, or antibody mixtures. In the case of antibody mixtures, 100 nM of each antibody was used. Following an additional wash, cells were resuspended in 5 ng/μl anti-human IgG-AlexaFluor647 (Jackson Immunoresearch 109-605-098) and incubated for 30 minutes at 4°C. Cells were washed once more and resuspended in 1 ng/μl anti-FLAG-PE before 30 minutes of incubation at 4°C. Cells were then pelleted by centrifugation at 8000 x g for 30s and kept on ice until sorting. Cells expressing RBD that maintained antibody-binding (IgG+/FLAG+) or showed a complete loss of antibody binding (escape) (IgG-/FLAG+) were sorted by FACS (BD Aria Fusion or Sony MA800 instrument). Collected cells were cultured in SD-UT medium for 16-40 hours at 30 °C. Induction and sorting was repeated for multiple rounds until the desired populations of RBD variants showed purity for binding and escape (non-binding) to antibodies.

### Antibody production and purification

Heavy chain and light chain inserts for REGN10933, REGN10987 (PDB: 6XDG) and LY-CoV16 (PDB: 7C01), LY-CoV555 (PDB: 7KMG) were cloned into pTwist transient expression vectors by Gibson Assembly. 30 mL cultures of Expi293 cells (Thermo, A14635) were transfected according to the manufacturer’s instructions. After 5-7 days, dense Expi293 cultures were centrifuged at 300 x g for 5 minutes to pellet the cells. Supernatant was filtered using Steriflip^®^ 0.22 μm (Merck, SCGP00525) filter units. Using protein G purification, Expi supernatant was directly loaded onto Protein G Agarose (Pierce, Cat# 20399) gravity columns, washed twice with PBS and eluted using Protein G Elution Buffer (Pierce, Cat# 21004). The eluted fractions were immediately neutralized with 1M TRIS-Buffer (pH 8) to physiological pH and quantified by Nanodrop™ 2000c for A280 nm absorption. Protein containing fractions were pooled and buffer exchanged using SnakeSkin™ dialysis tubing (10 MWCO, Pierce Cat#68100) followed by further dialysis and concentration using Amicon Ultra-4 10kDa centrifugal units (Merck, Cat# UFC801096), as described previously (*5*).

### Deep sequencing of RBD libraries

Plasmid DNA encoding the RBD variants was isolated following the manufacturer’s instructions (Zymo D2004). Mutagenized regions of the RBD were amplified using custom oligonucleotides. Illumina Nextera barcode sequences were added in a second PCR amplification step, allowing for multiplexed high-throughput sequencing runs. Populations were pooled at the desired ratios and sequenced using Illumina 2 x 250 PE or 2 x 150 PE protocols (MiSeq or NovaSeq instruments).

### Processing of deep sequencing data, statistical analysis and plots

#### Data preprocessing

Sequencing reads were paired, quality trimmed and assembled using Geneious and BBDuk, with a quality threshold of qphred ≥ 25. Mutagenized regions of interest were then extracted using custom Python scripts, followed by translation to amino acid sequences. The sequences obtained from each of the three libraries (2C, 2CE and 2T) were pre-processed separately before being combined into the final training set used for model training and evaluation. To remove sequencing errors, all libraries were filtered for sequences complying with the initial degenerate codon mutagenesis scheme. Library 2CE was filtered for only those sequences retaining unmutated residues in positions 417/439, to focus on the 484-505 region. Next, library 2T was filtered using a threshold of read counts > 4 and restricted to sequences that were ≤ ED_3_ from Wu-Hu-1 RBD sequence.

Duplicate sequences in the full dataset were removed and a balanced dataset was created from the remaining data such that equal numbers of positive (binding) and negative class (non-binding) sequences were present for each ED. We observed significant bias in model performance when predictions are separated by ED from the Wu-Hu-1 RBD sequence, which was likely due to class imbalance in the training data. Class balancing was thus performed through random subsampling from the majority class at each ED equal to the counts from the minority class. Those that were not sampled from the majority class were then reserved separately as additional “unseen sequences”. These were then used for model evaluation to ensure that the models could generalize well even to the sequences removed during balancing.

#### Statistical analysis and plots

Statistical analysis was performed using R 4.0.1 (*6*) and Python 3.8.5 (*7*). Graphics were generated using the ggplot2 3.3.3 (*8*), ComplexHeatmap 2.4.3 (*9*) pheatmap 1.0.12 (*10*), igraph 1.2.6 (*11*), RCy3 2.8.1 (*12*), stringr 1.4.0 (*13*), dplyr 1.0.6 (*14*), and RColorBrewer 1.1-2 (*15*) R package.

#### Escape Networks

Network plots were generated using the igraph package 1.2.6 (*11*) and Cytoscape software 3.8.2 (*16*) with edges drawn between every pair of two amino acid sequences from ED 1 and 2, when the pair of sequences share a common mutation on amino acid level. Edges were colored according to the change in number of antibodies that escape. Nodes representing RBD variant sequences were clustered and colored according to the number of antibodies that escape, and the mutational distance from the reference sequence.

### Machine learning model training and evaluation

All machine learning (ML) classifier models were built in Python (3.8.5) (*7*). Data was prepared and visualized using numpy (1.19.2), matplotlib (3.3.4), and pandas (1.2.4). Random Forest (RF) and other benchmarking ML models were built using Scikit-Learn (0.24.2), a 80/20 train-test data split (random split) to train baseline models, and a 90/10 traintest data split (random split) for final RF and RNN models. Keras libraries (2.4.3) from Tensorflow (v2.5) were used to build the long-short-term-memory recurrent neural networks (RNN) models.

RBD sequences were one-hot encoded prior to being used as inputs into the models. For the RNN, the 2D one-hot encoded matrix was used as the input, while for all other models, the matrix was flattened into a single dimensional vector beforehand. After selecting the best models, hyperparameter optimization was performed to further improve the performance of the chosen RF and RNN models using 50 rounds of Random Search with 5-fold cross-validation while scoring based on precision. All RF models were further calibrated using both the “isotonic” and Platt scaling (*17-19*), and the best model from the three was selected by calculating the overall mean-square error (MSE) from the true labels, with the RF model with the lowest MSE selected as the final model. For evaluation of the models on the held-out test set, models were evaluated on the basis of Accuracy, F1, Precision, Recall, and AUC-ROC curve using the entire test set. For further detailed evaluation, the test data was separated into two distances: low and high distance sequence sets, which consisted only of sequences ≤ ED_5_ or ≥ ED_6_ from Wu-Hu-1 RBD sequence, respectively. These two sets were then used to evaluate the accuracy, F1, Precision, and Recall of models to investigate any performance bias at different distances. The accuracy of the models were also evaluated on the sequences reserved during dataset balancing (‘unseen sequences’) separated similarly by ED.

### *In silico* sequence generation and evaluation

Synthetic RBD variant sequences were generated *in silico* using custom Python scripts for selected edit distances (ED) from the Wu-Hu-1 RBD sequence. The ED was defined on both the nucleotide and amino acid level, such that each generated nucleotide sequence was categorized by an ED pair (distance_nt, distance_aa). The synthetic variants (*in silico* generated sequences) were evaluated for their probability of ACE2-binding and non-binding using a consensus model (RF and RNN) approach. For a given RBD sequence, ACE2-binding prediction was defined as the case where both models output *P* > 0.5, else the sequence was considered as ACE2 non-binding. Similarly, the sequences were evaluated for binding and escape (non-binding) from monoclonal antibodies. Here the sequences were categorized into one of four categories: escape (both models *P* <0.25), antibody binding (both models *P* >0.75), unsure (at least one model gives *P* between 0.25 and 0.75), and disagree (one model outputs *P* <0.25 while the other model outputs a *P* >0.75).

### Experimental validation of selected RBD variants for ACE2-binding and antibody escape

Individual sequences for RBD variants were ordered as complementary forward and reverse primers (Integrated DNA Technologies) in 96-well plates A single round of annealing and extension was used to produce double-stranded DNA with 14-bp of homology at 5’ and 3’ ends to the pYD1-RBD entry vector, followed by Gibson Assembly with EcoRI digested vector. Plasmids were transformed into EBY100 prepared with the Frozen-EZ Yeast Transformation Kit II (Zymo) and plated on SD-UT agar. Individual colonies were picked and grown in SD-UT liquid medium overnight at 30°C, then diluted to OD_600_ = 0.5 in SG-UT medium and grown for 40-48 hours at 23°C. Cells were stained with biotinylated ACE2 or purified antibody as described above. Flow cytometry analysis was performed on the BD Fortessa cytometer.

### Structural Prediction of RBD variants by AlphaFold2

Structural predictions were generated with the Alphafold v2.1.0 public iPython notebook using residues 331-530 of the spike protein. (https://colab.research.google.com/github/deepmind/alphafold/blob/main/notebooks/AlphaFold.ipynb) (*20*). Results were visualized and aligned in PyMol v2.2.3 (*21*).

---

**Supplementary Figure 1.**
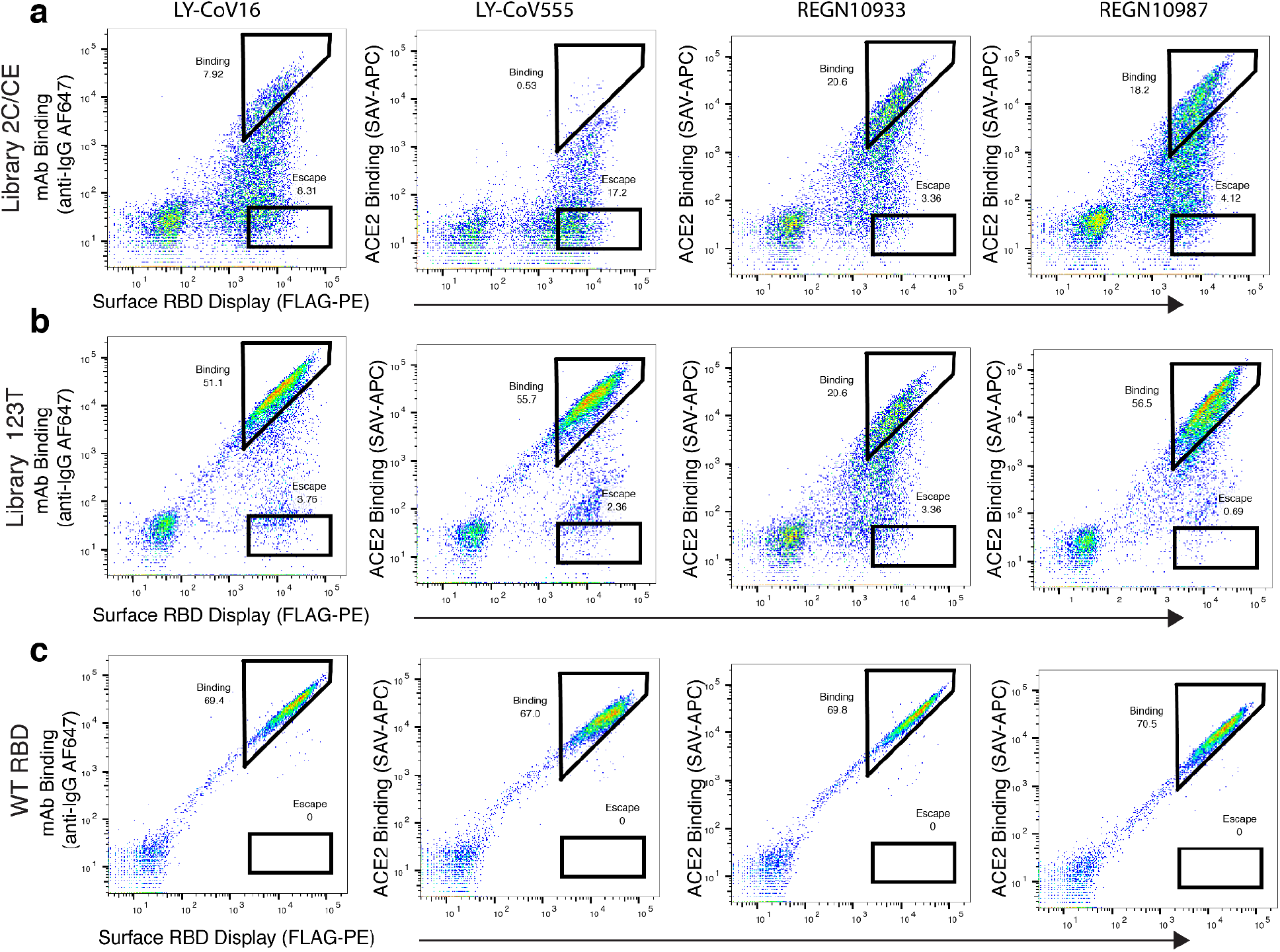
Selection of RBD variants for antibody binding and escape by flow cytometry. Yeast display of RBD libraries pre-selected for ACE2 binding were sorted by flow cytometry for binding and escape to four therapeutic monoclonal antibodies (mAbs): LY-CoV16, LY-CoV555, REGN10933, and REGN10987. (**a**) The RBD libraries 2C and 2CE were pooled together; (**b**) library 2T was pooled with libraries 1T and 3T. The original Wu-Hu-1 RBD (**c**) was used as a control for antibody binding and escape. Approximately 10^7^ yeast cells were screened for each antibody (**Supplementary Table 1**).

**Supplementary Figure 2.**
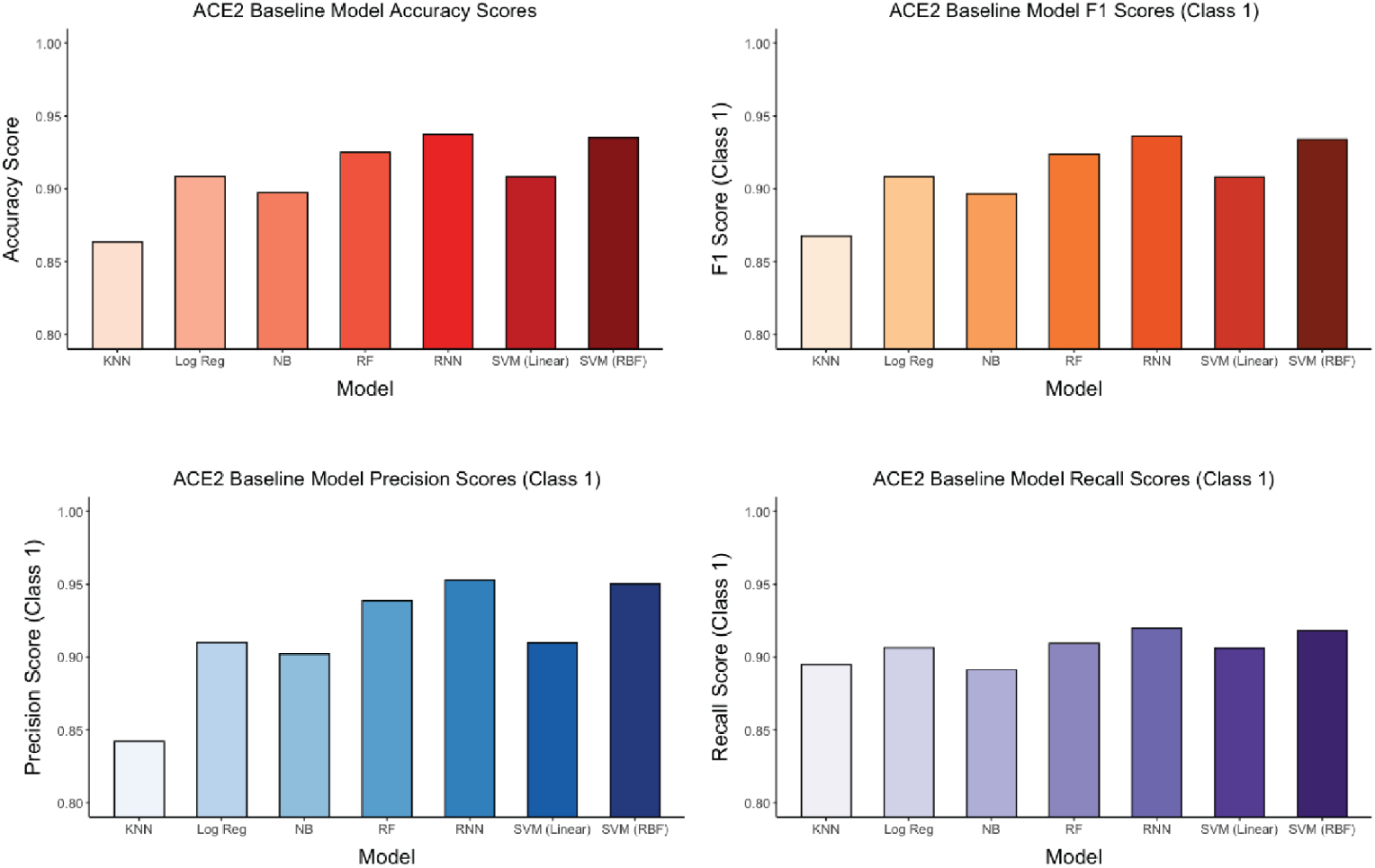
Performance metrics for seven different baseline machine learning models. K-nearest Neighbours (KNN), Logistic Regression (Log Reg), Naive Bayes (NB), Random Forest (RF), Long-short term memory recurrent neural network (RNN), Support vector machine with linear kernel (SVM Linear), and Support vector machine with radial basis function kernel (SVM RBF) models were trained on the ACE2 deep sequencing data without hyperparameter optimization. Models were then challenged to perform classification by predicting a probability (*P*) of ACE2 binding on test data. Performance of models was evaluated by Accuracy, F1, Precision, and Recall. All models except RNN were trained using Sci-kit Learn, and the RNN was trained using Keras.

**Supplementary Figure 3.**
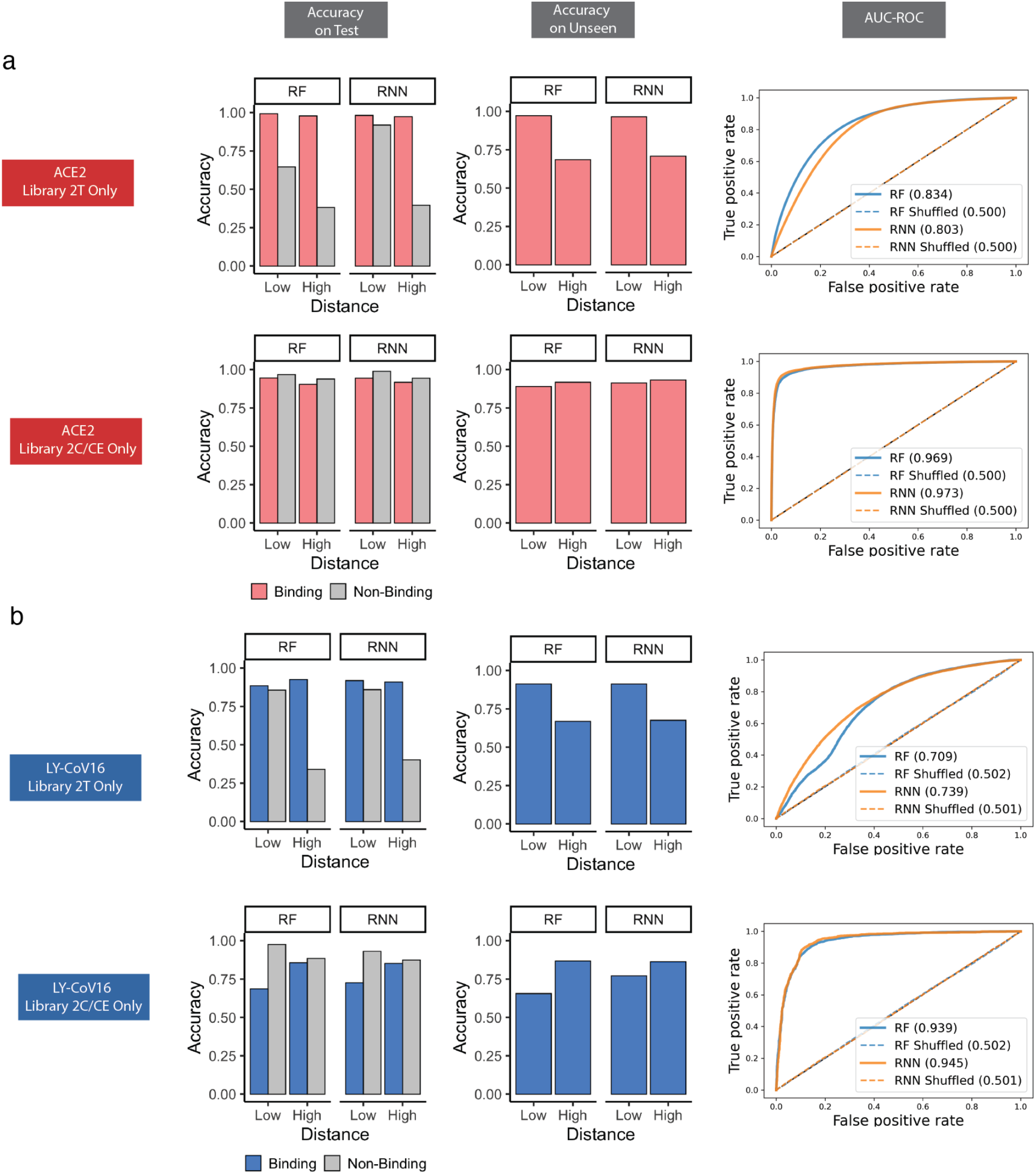
Accuracy and ROC curves for models trained on data from low and high distance libraries. **a, b,** The training data for ACE2 or LY-CoV16 were split into two subsets to train supervised machine learning (Random Forest, RF) and deep learning (recurrent neural network, RNN) models. Library 2T training data contained only sequences that were ≤ ED _3_ and Library 2C/CE training data contained only sequences that were ≥ ED_4_ from original Wu-Hu-1 RBD. The models then performed classification on the entirety of the held-out test data, and the unseen data were removed during balancing. Low and high distance sequences are defined as those ≤ ED_5_ and ≥ ED_6_; from Wu-Hu-1 RBD, respectively. Model performances are evaluated by accuracy and AUC-ROC.

**Supplementary Figure 4.**
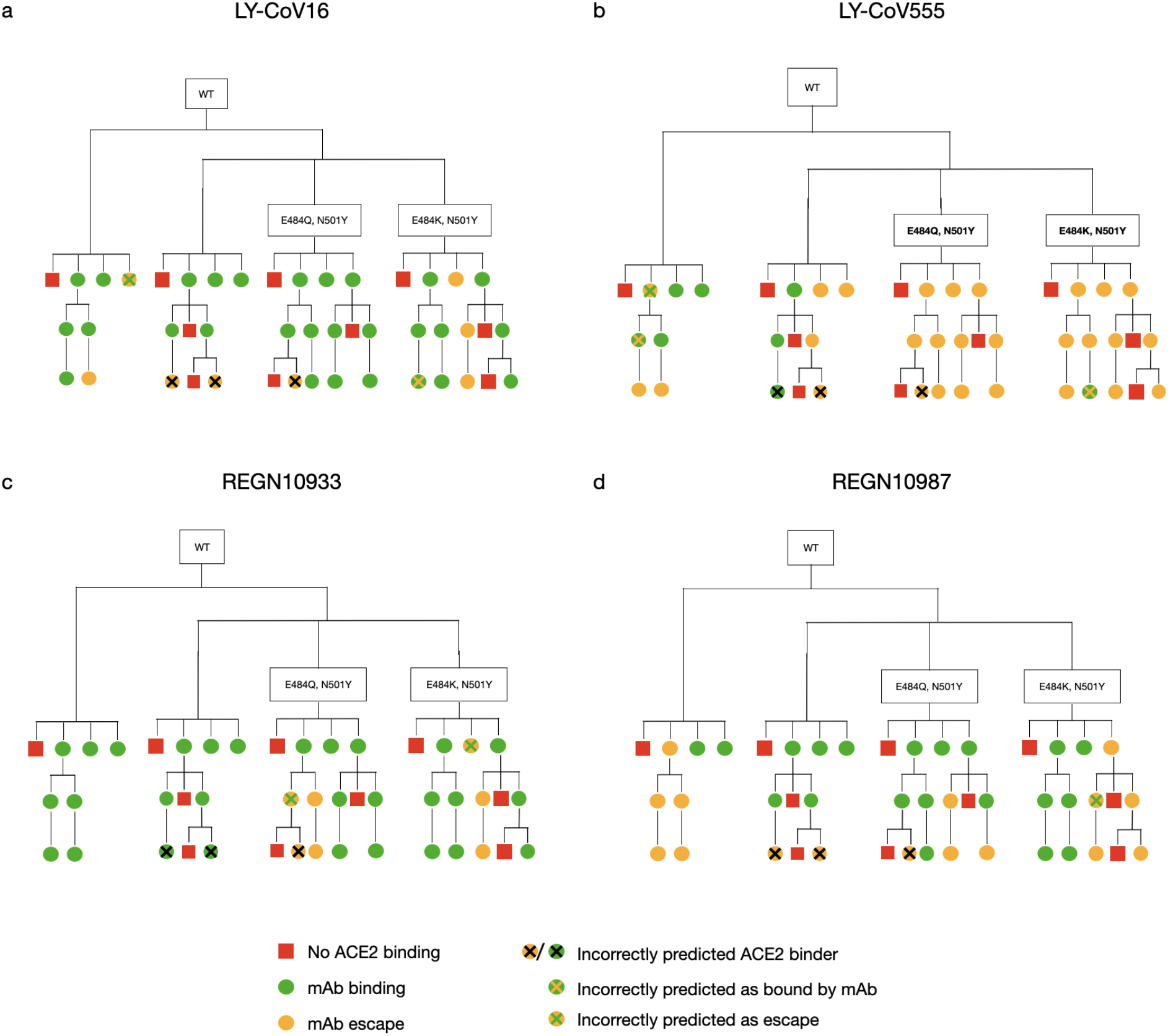
Evaluation of antibody escape predictions for synthetic lineages. **a-d,** RBD sequences at chosen EDs (ED_0_, ED_3_, ED_5_, ED_7_) from the Wu-Hu-1 RBD were predicted for ACE2 binding and escape from four therapeutic monoclonal antibodies (mAbs). Accuracy for antibody escape predictions are the following: LY-CoV16 = 31/33 (93.94%), LY-CoV555 = 30/33 (90.91%), REGN10933 = 31/33 (93.94%), REGN10987 = 32/33 (96.97%).

**Supplementary Figure 5.**
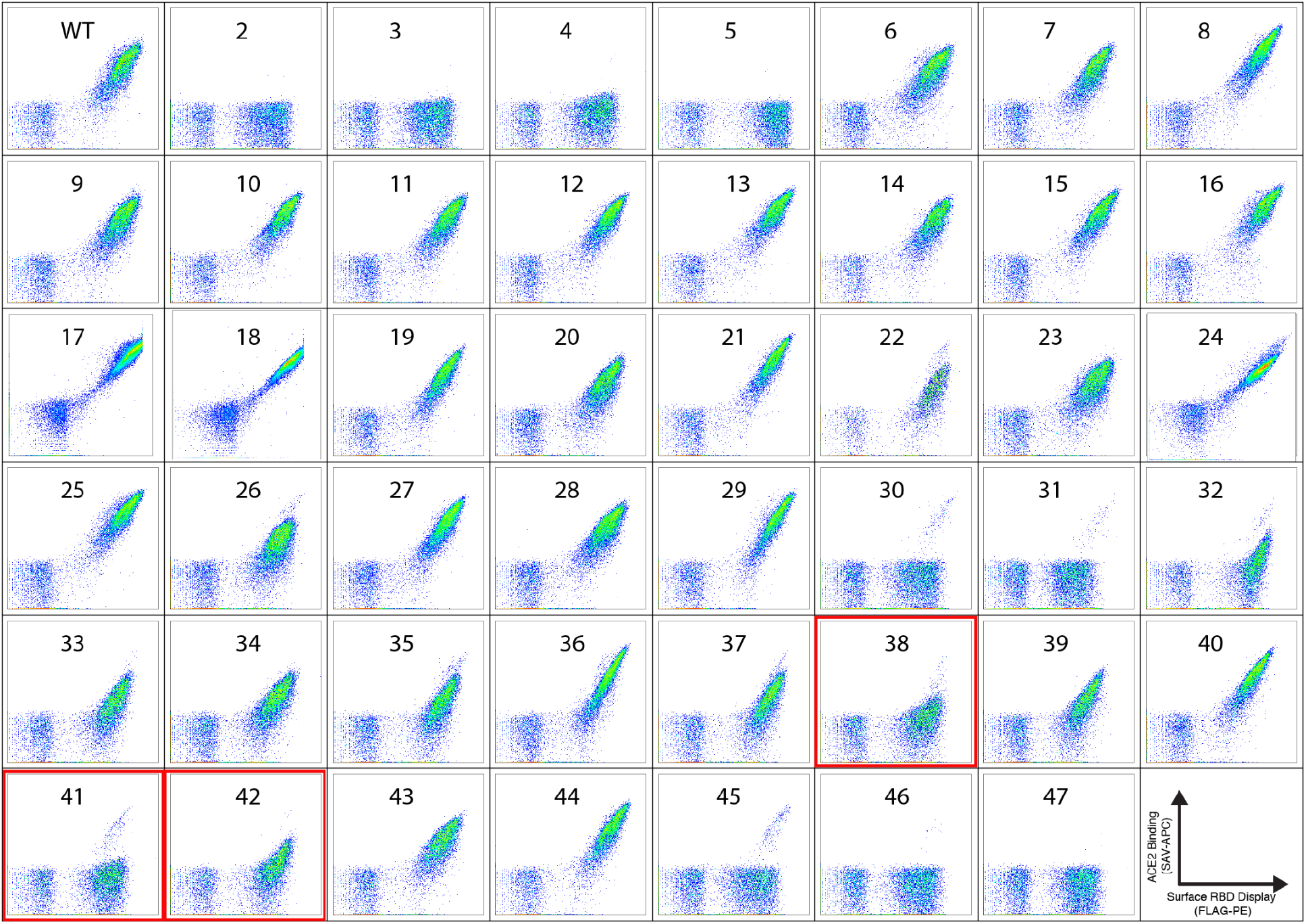
Yeast display screening of synthetic RBD variants for ACE2 binding. The 46 selected synthetic variants (see **Fig. 4**) were individually cloned and expressed for yeast display and ACE2 binding by flow cytometry. 43 variants showed ACE2 binding or non-binding that matched machine learning predictions. The ACE2-binding status for two variants (38 and 42) could not be conclusively determined, while one variant (41) showed was incorrectly predicted by machine learning for ACE2 binding.

**Supplementary Figure 6.**
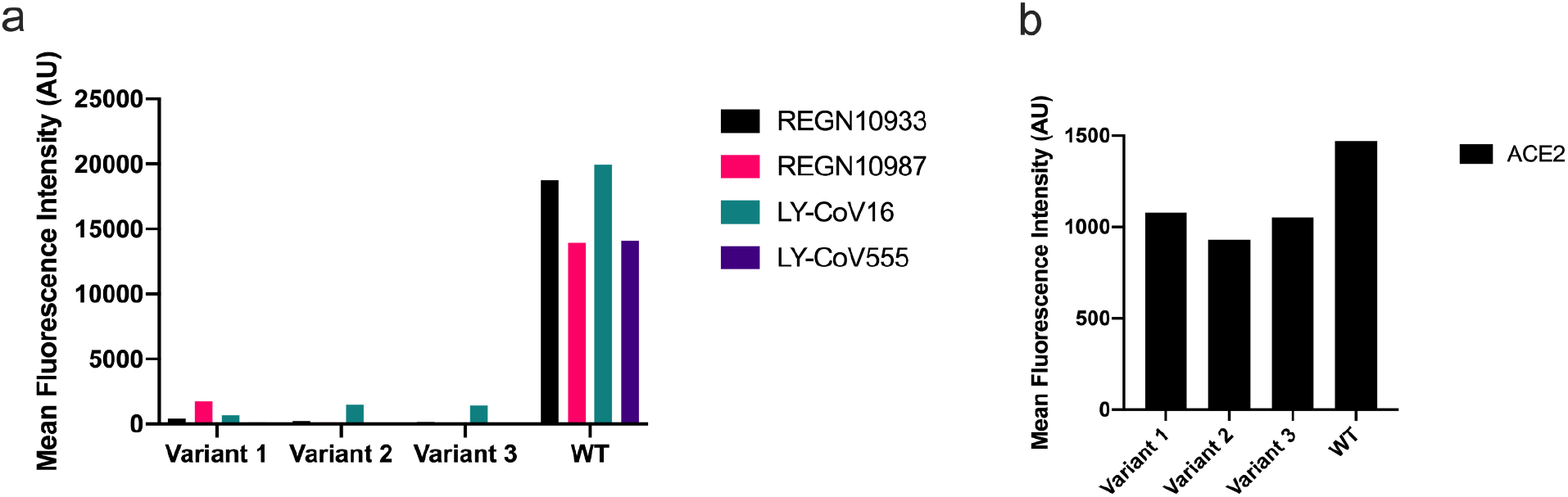
Experimental validation of synthetic RBD variants predicted to escape all four therapeutic antibodies. Three synthetic RBD variants of ED_3_ from Wu-Hu-1 RBD that were predicted to escape all four therapeutic antibodies by the consensus machine learning model were expressed as individual clones in yeast and evaluated by flow cytometry for binding to antibody (**a**) or ACE2 (**b**).

**Supplementary Figure 7.**
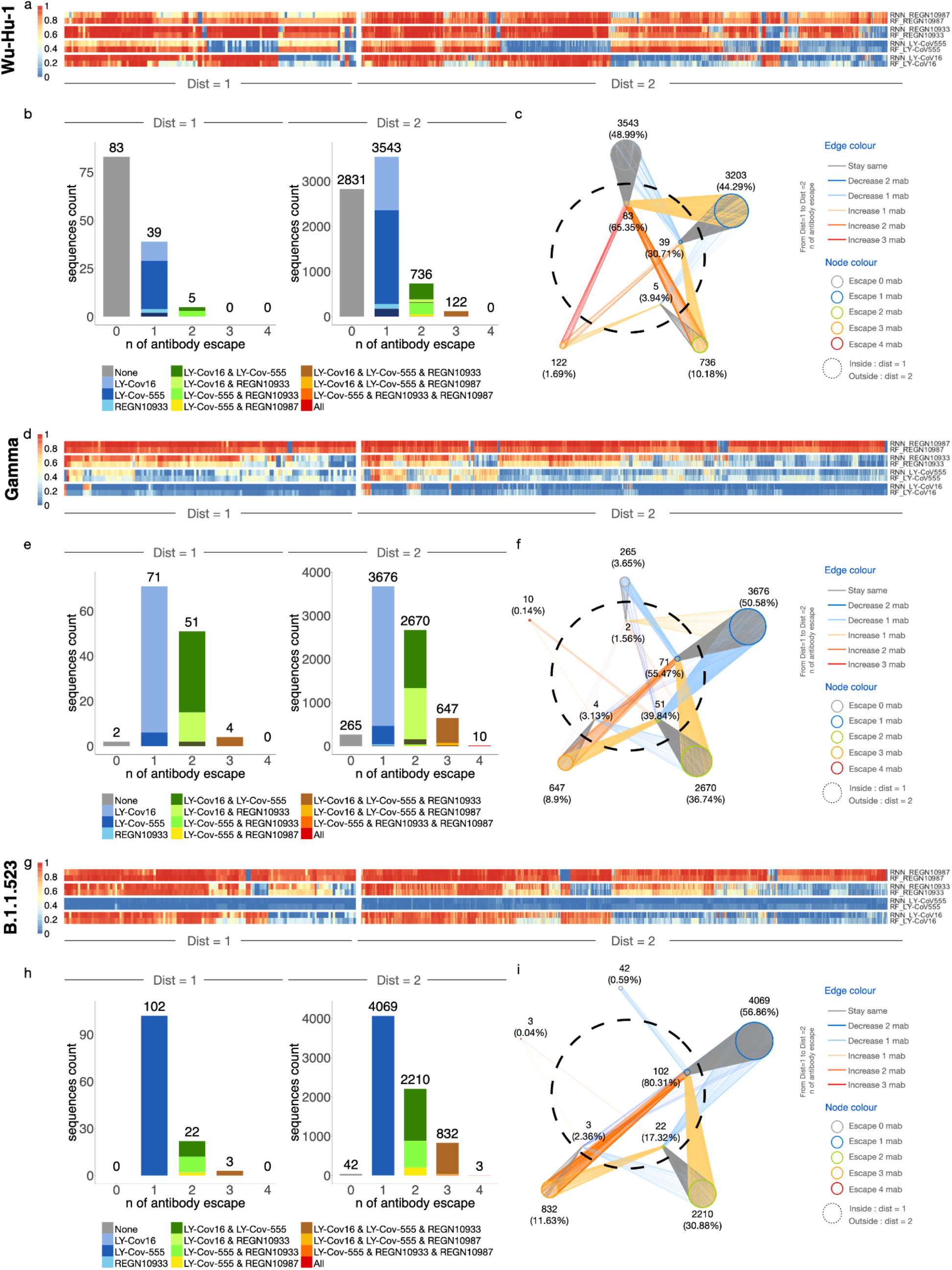
Predictive profiling of additional selected RBD variants for antibody escape across low mutational distances. **a,d,g,** Heatmap depicts monoclonal antibody (mAb) binding as assessed by RF and RNN models of ED_1_ and ED_2_ variants of Wu-Hu-1, gamma, and B.1.523. **b, e, h,** The number of sequences escaping a combination of *n* (number) mAbs for ED_1_ and ED_2_ (agreement between models, threshold >0.5). **c, f, i,** Deep escape networks display possible evolutionary paths between variants and their escape from mAbs.

**Supplementary Figure 8.**
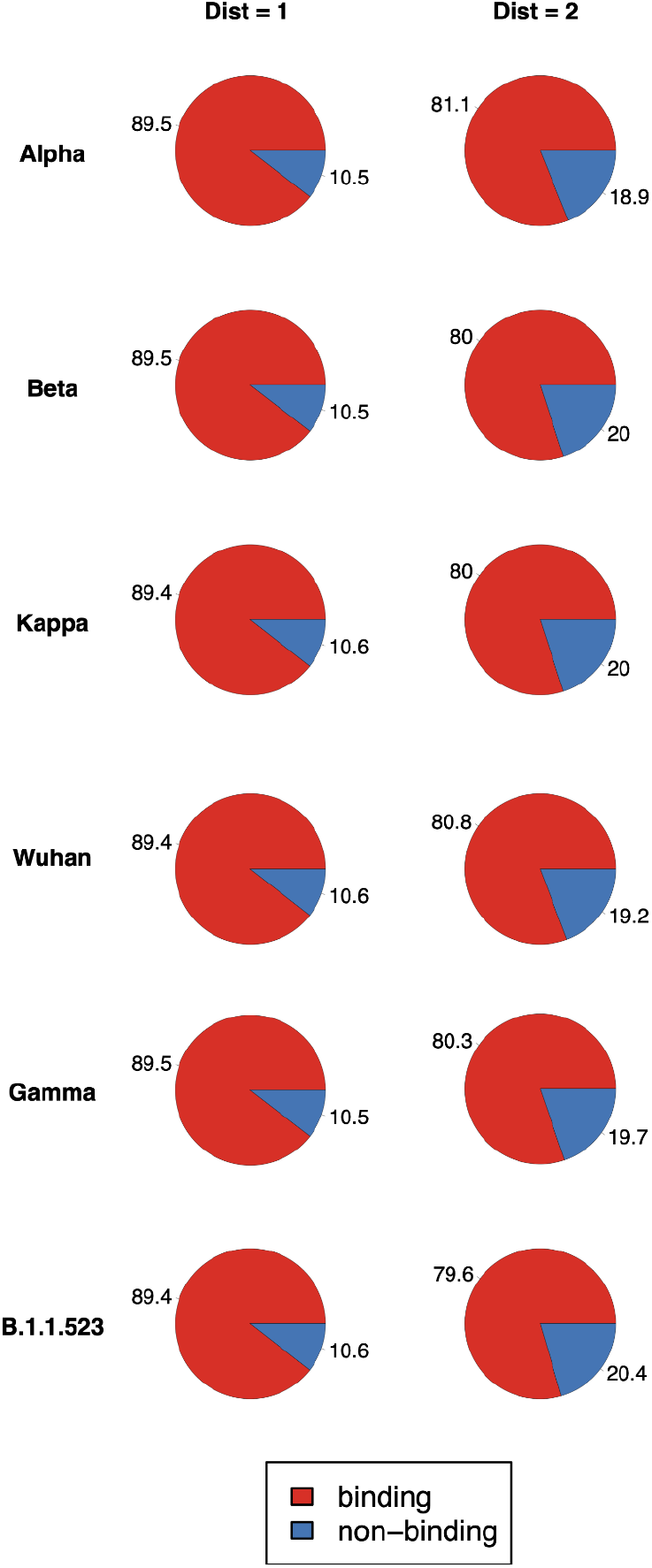
ACE2 binding predictions of synthetic RBD variants. Synthetic variants at ED_1_ and ED_2_ from alpha, beta, kappa, Wu-Hu-1, gamma B.1.1.523 were assessed for ACE2 binding using the RF and RNN models. Variants where both models output of *P* > 0.5 were predicted as ACE2 binding, all others as non-binding.

**Supplementary Figure 9.**
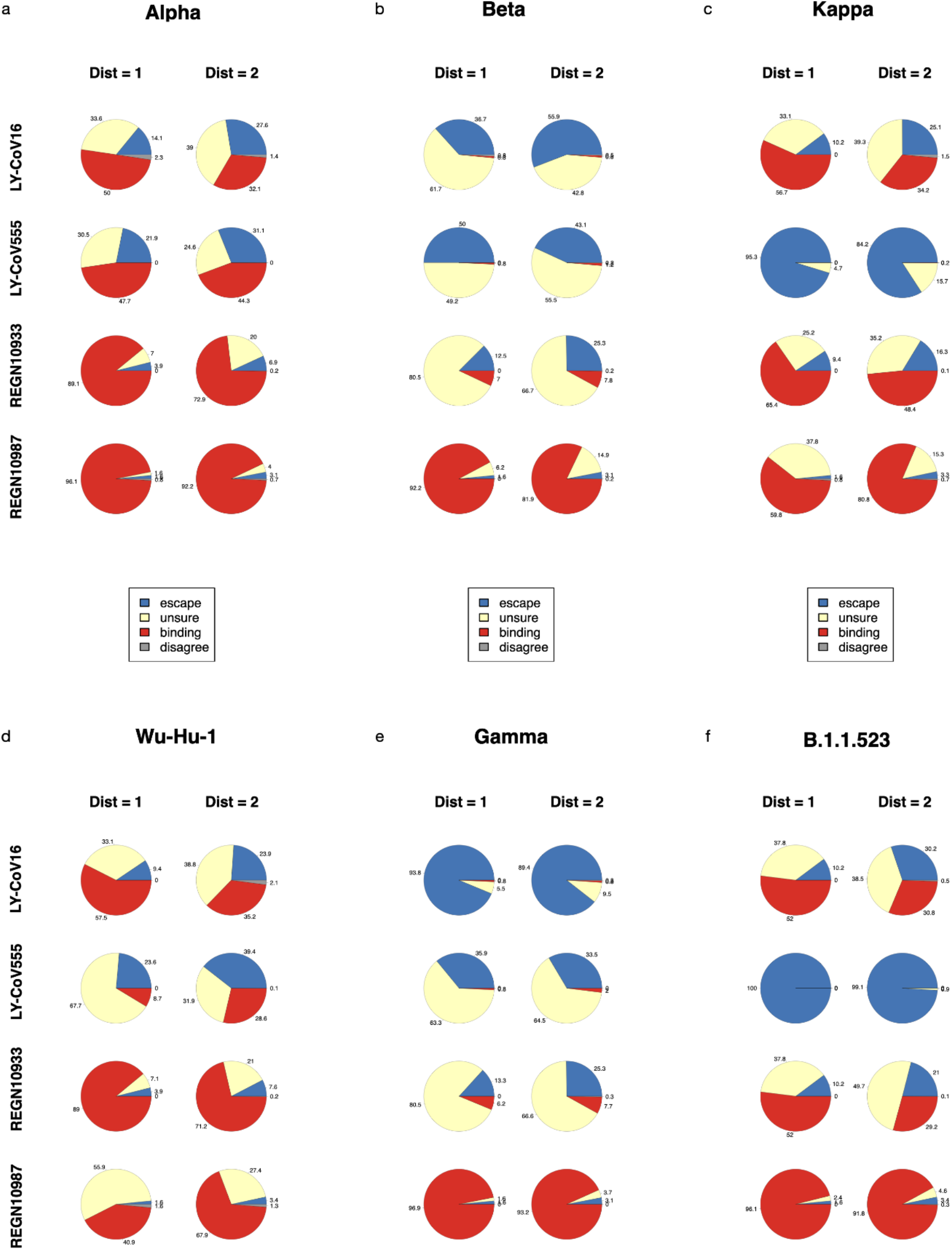
Antibody escape predictions of synthetic RBD variants. **a–f,** Synthetic RBD variants at edit acid distances 1 and 2 from alpha, beta, kappa, Wu-Hu-1, gamma, and B.1.1.523 were assessed for binding or escape to four therapeutic antibodies using the RF and RNN models. Variants where both models output a *P* > 0.25 were predicted as *escape*, when both models output *P* > 0.75 were predicted as *binding*, all other cases were determined to be *unsure* (at least one model with output of *P* between 0.25 and 0.75) or *disagree* (one mode output of *P*l < 0.25 and other model output of *P* > 0.75).

**Supplementary Table 1.** Sorting statistics for RBD library sorting.

| Population | Antigen | Events Screened | Binding Events Collected | Non-Binding Events Collected |
| --- | --- | --- | --- | --- |
| Library 2C | 50 nM ACE2 | 5.00E+07 | 1.60E+06 | 6.40E+06 |
| Library 2CE | 50 nM ACE2 | 5.00E+07 | 5.90E+05 | 6.40E+06 |
| Library 2C + 2CE, ACE2-binding | 100 nM REGN10933 | 1.07E+07 | 1.21E+06 | 3.93E+05 |
| Library 2C + 2CE, ACE2-binding | 100n nM RENG10987 | 1.02E+07 | 9.54E+05 | 5.41E+05 |
| Library 2C + 2CE, ACE2-binding | 100 nM Ly-CoV16 | 8.12E+06 | 5.07E+05 | 6.92E+05 |
| Library 2C + 2CE, ACE2-binding | 100 nM Ly-CoV555 | 1.13E+07 | 2.36E+04 | 1.87E+06 |
| Library 2T | 50 nM ACE2 | 1.52E+07 | 8.53E+05 | 1.49E+06 |
| Library 2T, ACE2-binding | 100 nM REGN10933 | 1.94E+06 | 5.09E+05 | 4.11E+04 |
| Library 2T, ACE2-binding | 100n nM RENG10987 | 2.43E+06 | 5.01E+05 | 4.57E+04 |
| Library 2T, ACE2-binding | 100 nM Ly-CoV16 | 2.56E+06 | 5.66E+05 | 1.13E+05 |
| Library 2T, ACE2-binding | 100 nM Ly-CoV555 | 3.15E+06 | 7.61E+05 | 1.48E+05 |

**Supplementary Table 2.** Sequencing statistics for RBD library sorting.

| Population | Antigen | Paired and Filtered Reads |  |
| --- | --- | --- | --- |
|  |  | Binding | Non-Binding |
| Library 2C | 50 nM ACE2 | 1.20E+06 | 1.16E+06 |
| Library 2CE | 50 nM ACE2 | 1.01E+06 | 1.48E+06 |
| Library 2C + 2CE, ACE2 binding | 100 nM REGN10933 | 9.54E+05 | 7.55E+05 |
| Library 2C + 2CE, ACE2 binding | 100n nM RENG10987 | 2.27E+05 | 1.15E+04 |
| Library 2C + 2CE, ACE2 binding | 100 nM Ly-CoV16 | 1.85E+06 | 2.98E+05 |
| Library 2C + 2CE, ACE2 binding | 100 nM Ly-CoV555 | 1.16E+05 | 5.47E+05 |
| Library 123T | 50 nM ACE2 | 1.53E+06 | 1.88E+06 |
| Library 123T, ACE2 binding | 100 nM REGN10933 | 5.53E+06 | 2.55E+06 |
| Library 123T, ACE2 binding | 100n nM RENG10987 | 3.30E+06 | 2.87E+06 |
| Library 123T, ACE2 binding | 100 nM Ly-CoV16 | 1.59E+06 | 3.20E+06 |
| Library 123T, ACE2 binding | 100 nM Ly-CoV555 | 3.64E+06 | 2.56E+06 |

**Supplementary Table 3 (see Excel File). Detailed counts of sequences from each dataset separated by ED from Wu-Hu-1 RBD and class label.** Counts include raw sequences, after balancing, and train/test split.

**Supplementary Table 4 (see Excel File). Detailed sequences used as the training data for individual models.** Each dataset combines sequences from 2T/2C/2CE libraries after preprocessing, filtering, and removing duplicates. All files are separated into train/test/unseen sequences.

**Supplementary Table 5.** Hyperparameter tuning values used for Random Forest and RNN Models. Hyperparameters were tuned though RandomSearchCV, scored for “precision”, using Sci-kit Learn for 50 rounds.

| Random Forest | RNN |
| --- | --- |
| n_estimators = 500,<br>min_samples_split = 2,<br>min_samples_leaf = 1,<br>max_depth = 150,<br>max_features = 'sqrt',<br>criterion = gini<br>HP tuning parameters:<br>n_estimators: [50-500]<br>min_samples_split: [2,5,10]<br>min_samples_leaf: [1,2,5]<br>max_features: ['auto', 'sqrt']<br>max_depth: [20-150,None] | 3 sequential LSTM/Dropout layers<br>Dense layer 50 units<br>Sigmoid activation output<br>Cross-entropy loss<br>HP tuning parameters:<br>batch size: [16, 32]<br>epochs: [10, 20]<br>dropout rate: [0.1,0.2],<br>LSTM units: [40, 80],<br>optimizer: [adam, rmsprop] |

**Supplementary Table 6 (see Excel File). Detailed metrics of all baseline models trained on ACE2 sequences.** Metrics evaluated include: total accuracy and F1, precision, and recall for both classes. Models were evaluated on the entirety of the held-out test set, without hyperparameter optimization.

**Supplementary Table 7 (see Excel File). Detailed metrics of all final models.** Metrics evaluated include: total accuracy and F1, precision, and recall for the positive class. Models were evaluated on the entirety of the held-out test set, or “Low Distance” and “High Distance” subsets of the held-out test set.

**Supplementary Table 8 (see Excel File). Machine and deep learning model predictions compared to susceptibility data from the Stanford Database (https://covdb.stanford.edu/page/susceptibility-data/, 2021-10-19).** RF and RNN model predictions are compared to previously published susceptibility data. The sequences had previously been excluded from the training datasets.

**Supplementary Table 9 (see Excel File). DML predictions on *in silico* generated sequences ED_1-2_ from selected variants.** RF and RNN model predictions.

## REFERENCES

1. P. Wang, M. S. Nair, L. Liu, S. Iketani, Y. Luo, Y. Guo, M. Wang, J. Yu, B. Zhang, P. D. Kwong, B. S. Graham, J. R. Mascola, J. Y. Chang, M. T. Yin, M. Sobieszczyk, C. A. Kyratsous, L. Shapiro, Z. Sheng, Y. Huang, D. D. Ho, Antibody resistance of SARS-CoV-2 variants B.1.351 and B.1.1.7. Nature. 593, 130–135 (2021).

2. M. Hoffmann, P. Arora, R. Groß, A. Seidel, B. F. Hörnich, A. S. Hahn, N. Krüger, L. Graichen, H. Hofmann-Winkler, A. Kempf, M. S. Winkler, S. Schulz, H.-M. Jäck, B. Jahrsdörfer, H. Schrezenmeier, M. Müller, A. Kleger, J. Münch, S. Pöhlmann, SARS-CoV-2 variants B.1.351 and P.1 escape from neutralizing antibodies. Cell. 184, 2384–2393.e12 (2021).

3. C. K. Wibmer, F. Ayres, T. Hermanus, M. Madzivhandila, P. Kgagudi, B. Oosthuysen, B. E. Lambson, T. de Oliveira, M. Vermeulen, K. van der Berg, T. Rossouw, M. Boswell, V. Ueckermann, S. Meiring, A. von Gottberg, C. Cohen, L. Morris, J. N. Bhiman, P. L. Moore, SARS-CoV-2 501Y.V2 escapes neutralization by South African COVID-19 donor plasma. Nat. Med. 27, 622–625 (2021).

4. W. F. Garcia-Beltran, E. C. Lam, K. S. Denis, A. D. Nitido, Z. H. Garcia, B. M. Hauser, J. Feldman, M. N. Pavlovic, D. J. Gregory, M. C. Poznansky, A. Sigal, A. G. Schmidt, A. J. Iafrate, V. Naranbhai, A. B. Balazs, Multiple SARS-CoV-2 variants escape neutralization by vaccine-induced humoral immunity. Cell. 184, 2372–2383.e9 (2021).

5. A. Fontanet, B. Autran, B. Lina, M. P. Kieny, S. S. A. Karim, D. Sridhar, SARS-CoV-2 variants and ending the COVID-19 pandemic. The Lancet. 397, 952–954 (2021).

6. F. Wu, S. Zhao, B. Yu, Y.-M. Chen, W. Wang, Z.-G. Song, Y. Hu, Z.-W. Tao, J.-H. Tian, Y.-Y. Pei, M.-L. Yuan, Y.-L. Zhang, F.-H. Dai, Y. Liu, Q.-M. Wang, J.-J. Zheng, L. Xu, E. C. Holmes, Y.-Z. Zhang, A new coronavirus associated with human respiratory disease in China. Nature. 579, 265–269 (2020).

7. K. D. McCormick, J. L. Jacobs, J. W. Mellors, The emerging plasticity of SARS-CoV-2. Science. 371, 1306–1308 (2021).

8. R. Yan, Y. Zhang, Y. Li, L. Xia, Y. Guo, Q. Zhou, Structural basis for the recognition of SARS-CoV-2 by full-length human ACE2. Science. 367, 1444–1448 (2020).

9. D. Zhou, W. Dejnirattisai, P. Supasa, C. Liu, A. J. Mentzer, H. M. Ginn, Y. Zhao, H. M. E. Duyvesteyn, A. Tuekprakhon, R. Nutalai, B. Wang, G. C. Paesen, C. Lopez-Camacho, J. Slon-Campos, B. Hallis, N. Coombes, K. Bewley, S. Charlton, T. S. Walter, D. Skelly, S. F. Lumley, C. Dold, R. Levin, T. Dong, A. J. Pollard, J. C. Knight, D. Crook, T. Lambe, E. Clutterbuck, S. Bibi, A. Flaxman, M. Bittaye, S. Belij-Rammerstorfer, S. Gilbert, W. James, M. W. Carroll, P. Klenerman, E. Barnes, S. J. Dunachie, E. E. Fry, J. Mongkolsapaya, J. Ren, D. I. Stuart, G. R. Screaton, Evidence of escape of SARS-CoV-2 variant B.1.351 from natural and vaccine-induced sera. Cell. 184, 2348–2361.e6 (2021).

10. C. Yi, X. Sun, J. Ye, L. Ding, M. Liu, Z. Yang, X. Lu, Y. Zhang, L. Ma, W. Gu, A. Qu, J. Xu, Z. Shi, Z. Ling, B. Sun, Key residues of the receptor binding motif in the spike protein of SARS-CoV-2 that interact with ACE2 and neutralizing antibodies. Cell. Mol. Immunol. 17, 621–630 (2020).

11. P. Supasa, D. Zhou, W. Dejnirattisai, C. Liu, A. J. Mentzer, H. M. Ginn, Y. Zhao, H. M. E. Duyvesteyn, R. Nutalai, A. Tuekprakhon, B. Wang, G. C. Paesen, J. Slon-Campos, C. López-Camacho, B. Hallis, N. Coombes, K. R. Bewley, S. Charlton, T. S. Walter, E. Barnes, S. J. Dunachie, D. Skelly, S. F. Lumley, N. Baker, I. Shaik, H. E. Humphries, K. Godwin, N. Gent, A. Sienkiewicz, C. Dold, R. Levin, T. Dong, A. J. Pollard, J. C. Knight, P. Klenerman, D. Crook, T. Lambe, E. Clutterbuck, S. Bibi, A. Flaxman, M. Bittaye, S. Belij-Rammerstorfer, S. Gilbert, D. R. Hall, M. A. Williams, N. G. Paterson, W. James, M. W. Carroll, E. E. Fry, J. Mongkolsapaya, J. Ren, D. I. Stuart, G. R. Screaton, Reduced neutralization of SARS-CoV-2 B.1.1.7 variant by convalescent and vaccine sera. Cell. 184, 2201–2211.e7 (2021).

12. P. Han, C. Su, Y. Zhang, C. Bai, A. Zheng, C. Qiao, Q. Wang, S. Niu, Q. Chen, Y. Zhang, W. Li, H. Liao, J. Li, Z. Zhang, H. Cho, M. Yang, X. Rong, Y. Hu, N. Huang, J. Yan, Q. Wang, X. Zhao, G. F. Gao, J. Qi, Molecular insights into receptor binding of recent emerging SARS-CoV-2 variants. Nat. Commun. 12, 6103 (2021).

13. S. C. A. Nielsen, F. Yang, K. J. L. Jackson, R. A. Hoh, K. Röltgen, G. H. Jean, B. A. Stevens, J.-Y. Lee, A. Rustagi, A. J. Rogers, A. E. Powell, M. Hunter, J. Najeeb, A. R. Otrelo-Cardoso, K. E. Yost, B. Daniel, K. C. Nadeau, H. Y. Chang, A. T. Satpathy, T. S. Jardetzky, P. S. Kim, T. T. Wang, B. A. Pinsky, C. A. Blish, S. D. Boyd, Human B Cell Clonal Expansion and Convergent Antibody Responses to SARS-CoV-2. Cell Host Microbe. 28, 516–525.e5 (2020).

14. F. Yang, S. C. A. Nielsen, R. A. Hoh, K. Röltgen, O. F. Wirz, E. Haraguchi, G. H. Jean, J.-Y. Lee, T. D. Pham, K. J. L. Jackson, K. M. Roskin, Y. Liu, K. Nguyen, R. S. Ohgami, E. M. Osborne, K. C. Nadeau, C. U. Niemann, J. Parsonnet, S. D. Boyd, Shared B cell memory to coronaviruses and other pathogens varies in human age groups and tissues. Science. 372, 738–741 (2021).

15. S. J. Zost, P. Gilchuk, R. E. Chen, J. B. Case, J. X. Reidy, A. Trivette, R. S. Nargi, R. E. Sutton, N. Suryadevara, E. C. Chen, E. Binshtein, S. Shrihari, M. Ostrowski, H. Y. Chu, J. E. Didier, K. W. MacRenaris, T. Jones, S. Day, L. Myers, F. Eun-Hyung Lee, D. C. Nguyen, I. Sanz, D. R. Martinez, P. W. Rothlauf, L.-M. Bloyet, S. P. J. Whelan, R. S. Baric, L. B. Thackray, M. S. Diamond, R. H. Carnahan, J. E. Crowe, Rapid isolation and profiling of a diverse panel of human monoclonal antibodies targeting the SARS-CoV-2 spike protein. Nat. Med. 26, 1422–1427 (2020).

16. C. O. Barnes, A. P. West, K. E. Huey-Tubman, M. A. G. Hoffmann, N. G. Sharaf, P. R. Hoffman, N. Koranda, H. B. Gristick, C. Gaebler, F. Muecksch, J. C. C. Lorenzi, S. Finkin, T. Hägglöf, A. Hurley, K. G. Millard, Y. Weisblum, F. Schmidt, T. Hatziioannou, P. D. Bieniasz, M. Caskey, D. F. Robbiani, M. C. Nussenzweig, P. J. Bjorkman, Structures of human antibodies bound to SARS-CoV-2 spike reveal common epitopes and recurrent features of antibodies. Cell. 0 (2020), doi:10.1016/j.cell.2020.06.025.

17. W. Dejnirattisai, D. Zhou, H. M. Ginn, H. M. E. Duyvesteyn, P. Supasa, J. B. Case, Y. Zhao, T. S. Walter, A. J. Mentzer, C. Liu, B. Wang, G. C. Paesen, J. Slon-Campos, C. López-Camacho, N. M. Kafai, A. L. Bailey, R. E. Chen, B. Ying, C. Thompson, J. Bolton, A. Fyfe, S. Gupta, T. K. Tan, J. Gilbert-Jaramillo, W. James, M. Knight, M. W. Carroll, D. Skelly, C. Dold, Y. Peng, R. Levin, T. Dong, A. J. Pollard, J. C. Knight, P. Klenerman, N. Temperton, D. R. Hall, M. A. Williams, N. G. Paterson, F. K. R. Bertram, C. A. Siebert, D. K. Clare, A. Howe, J. Radecke, Y. Song, A. R. Townsend, K.-Y. A. Huang, E. E. Fry, J. Mongkolsapaya, M. S. Diamond, J. Ren, D. I. Stuart, G. R. Screaton, The antigenic anatomy of SARS-CoV-2 receptor binding domain. Cell. 184, 2183–2200.e22 (2021).

18. W. T. Harvey, A. M. Carabelli, B. Jackson, R. K. Gupta, E. C. Thomson, E. M. Harrison, C. Ludden, R. Reeve, A. Rambaut, S. J. Peacock, D. L. Robertson, SARS-CoV-2 variants, spike mutations and immune escape. Nat. Rev. Microbiol., 1–16 (2021).

19. J. Hansen, A. Baum, K. E. Pascal, V. Russo, S. Giordano, E. Wloga, B. O. Fulton, Y. Yan, K. Koon, K. Patel, K. M. Chung, A. Hermann, E. Ullman, J. Cruz, A. Rafique, T. Huang, J. Fairhurst, C. Libertiny, M. Malbec, W.-Y. Lee, R. Welsh, G. Farr, S. Pennington, D. Deshpande, J. Cheng, A. Watty, P. Bouffard, R. Babb, N. Levenkova, C. Chen, B. Zhang, A. Romero Hernandez, K. Saotome, Y. Zhou, M. Franklin, S. Sivapalasingam, D. C. Lye, S. Weston, J. Logue, R. Haupt, M. Frieman, G. Chen, W. Olson, A. J. Murphy, N. Stahl, G. D. Yancopoulos, C. A. Kyratsous, Studies in humanized mice and convalescent humans yield a SARS-CoV-2 antibody cocktail. Science. 369, 1010–1014 (2020).

20. A. Baum, B. O. Fulton, E. Wloga, R. Copin, K. E. Pascal, V. Russo, S. Giordano, K. Lanza, N. Negron, M. Ni, Y. Wei, G. S. Atwal, A. J. Murphy, N. Stahl, G. D. Yancopoulos, C. A. Kyratsous, Antibody cocktail to SARS-CoV-2 spike protein prevents rapid mutational escape seen with individual antibodies. Science (2020), doi:10.1126/science.abd0831.

21. R. Shi, C. Shan, X. Duan, Z. Chen, P. Liu, J. Song, T. Song, X. Bi, C. Han, L. Wu, G. Gao, X. Hu, Y. Zhang, Z. Tong, W. Huang, W. J. Liu, G. Wu, B. Zhang, L. Wang, J. Qi, H. Feng, F.-S. Wang, Q. Wang, G. F. Gao, Z. Yuan, J. Yan, A human neutralizing antibody targets the receptor-binding site of SARS-CoV-2. Nature. 584, 120–124 (2020).

22. T. N. Starr, A. J. Greaney, A. S. Dingens, J. D. Bloom, Complete map of SARS-CoV-2 RBD mutations that escape the monoclonal antibody LY-CoV555 and its cocktail with LY-CoV016. Cell Rep. Med. 0 (2021), doi:10.1016/j.xcrm.2021.100255.

23. D. Pinto, Y.-J. Park, M. Beltramello, A. C. Walls, M. A. Tortorici, S. Bianchi, S. Jaconi, K. Culap, F. Zatta, A. De Marco, A. Peter, B. Guarino, R. Spreafico, E. Cameroni, J. B. Case, R. E. Chen, C. Havenar-Daughton, G. Snell, A. Telenti, H. W. Virgin, A. Lanzavecchia, M. S. Diamond, K. Fink, D. Veesler, D. Corti, Cross-neutralization of SARS-CoV-2 by a human monoclonal SARS-CoV antibody. Nature, 1–10 (2020).

24. P. L. Tzou, K. Tao, J. Nouhin, S.-Y. Rhee, B. D. Hu, S. Pai, N. Parkin, R. W. Shafer, Coronavirus Antiviral Research Database (CoV-RDB): An Online Database Designed to Facilitate Comparisons between Candidate Anti-Coronavirus Compounds. Viruses. 12, 1006 (2020).

25. J. ter Meulen, E. N. van den Brink, L. L. M. Poon, W. E. Marissen, C. S. W. Leung, F. Cox, C. Y. Cheung, A. Q. Bakker, J. A. Bogaards, E. van Deventer, W. Preiser, H. W. Doerr, V. T. Chow, J. de Kruif, J. S. M. Peiris, J. Goudsmit, Human monoclonal antibody combination against SARS coronavirus: synergy and coverage of escape mutants. PLoS Med. 3, e237 (2006).

26. M. Yuan, N. C. Wu, X. Zhu, C.-C. D. Lee, R. T. Y. So, H. Lv, C. K. P. Mok, I. A. Wilson, A highly conserved cryptic epitope in the receptor binding domains of SARS-CoV-2 and SARS-CoV. Science. 368, 630–633 (2020).

27. Y. Wu, F. Wang, C. Shen, W. Peng, D. Li, C. Zhao, Z. Li, S. Li, Y. Bi, Y. Yang, Y. Gong, H. Xiao, Z. Fan, S. Tan, G. Wu, W. Tan, X. Lu, C. Fan, Q. Wang, Y. Liu, C. Zhang, J. Qi, G. F. Gao, F. Gao, L. Liu, A noncompeting pair of human neutralizing antibodies block COVID-19 virus binding to its receptor ACE2. Science (2020), doi:10.1126/science.abc2241.

28. D. M. Fowler, S. Fields, Deep mutational scanning: a new style of protein science. Nat. Methods. 11, 801–807 (2014).

29. T. N. Starr, A. J. Greaney, S. K. Hilton, D. Ellis, K. H. D. Crawford, A. S. Dingens, M. J. Navarro, J. E. Bowen, M. A. Tortorici, A. C. Walls, N. P. King, D. Veesler, J. D. Bloom, Deep Mutational Scanning of SARS-CoV-2 Receptor Binding Domain Reveals Constraints on Folding and ACE2 Binding. Cell. 182, 1295–1310.e20 (2020).

30. T. N. Starr, A. J. Greaney, A. Addetia, W. W. Hannon, M. C. Choudhary, A. S. Dingens, J. Z. Li, J. D. Bloom, Prospective mapping of viral mutations that escape antibodies used to treat COVID-19. Science. 371, 850–854 (2021).

31. A. J. Greaney, T. N. Starr, P. Gilchuk, S. J. Zost, E. Binshtein, A. N. Loes, S. K. Hilton, J. Huddleston, R. Eguia, K. H. D. Crawford, A. S. Dingens, R. S. Nargi, R. E. Sutton, N. Suryadevara, P. W. Rothlauf, Z. Liu, S. P. J. Whelan, R. H. Carnahan, J. E. Crowe, J. D. Bloom, Complete Mapping of Mutations to the SARS-CoV-2 Spike Receptor-Binding Domain that Escape Antibody Recognition. Cell Host Microbe. 29, 44–57.e9 (2021).

32. A. J. Greaney, A. N. Loes, K. H. D. Crawford, T. N. Starr, K. D. Malone, H. Y. Chu, J. D. Bloom, Comprehensive mapping of mutations in the SARS-CoV-2 receptor-binding domain that affect recognition by polyclonal human plasma antibodies. Cell Host Microbe. 29, 463–476.e6 (2021).

33. J. Lan, J. Ge, J. Yu, S. Shan, H. Zhou, S. Fan, Q. Zhang, X. Shi, Q. Wang, L. Zhang, X. Wang, Structure of the SARS-CoV-2 spike receptor-binding domain bound to the ACE2 receptor. Nature. 581, 215–220 (2020).

34. D. M. Mason, C. R. Weber, C. Parola, S. M. Meng, V. Greiff, W. J. Kelton, S. T. Reddy, High-throughput antibody engineering in mammalian cells by CRISPR/Cas9-mediated homology-directed mutagenesis. Nucleic Acids Res. (2018), doi:10.1093/nar/gky550.

35. E. C. Thomson, L. E. Rosen, J. G. Shepherd, R. Spreafico, A. da Silva Filipe, J. A. Wojcechowskyj, C. Davis, L. Piccoli, D. J. Pascall, J. Dillen, S. Lytras, N. Czudnochowski, R. Shah, M. Meury, N. Jesudason, A. De Marco, K. Li, J. Bassi, A. O’Toole, D. Pinto, R. M. Colquhoun, K. Culap, B. Jackson, F. Zatta, A. Rambaut, S. Jaconi, V. B. Sreenu, J. Nix, I. Zhang, R. F. Jarrett, W. G. Glass, M. Beltramello, K. Nomikou, M. Pizzuto, L. Tong, E. Cameroni, T. I. Croll, N. Johnson, J. Di Iulio, A. Wickenhagen, A. Ceschi, A. M. Harbison, D. Mair, P. Ferrari, K. Smollett, F. Sallusto, S. Carmichael, C. Garzoni, J. Nichols, M. Galli, J. Hughes, A. Riva, A. Ho, M. Schiuma, M. G. Semple, P. J. M. Openshaw, E. Fadda, J. K. Baillie, J. D. Chodera, S. J. Rihn, S. J. Lycett, H. W. Virgin, A. Telenti, D. Corti, D. L. Robertson, G. Snell, Circulating SARS-CoV-2 spike N439K variants maintain fitness while evading antibody-mediated immunity. Cell. 184, 1171–1187.e20 (2021).

36. K.-C. Tsai, Y.-C. Lee, T.-S. Tseng, Comprehensive Deep Mutational Scanning Reveals the Immune-Escaping Hotspots of SARS-CoV-2 Receptor-Binding Domain Targeting Neutralizing Antibodies. Front. Microbiol. 12, 1812 (2021).

37. E. T. Boder, K. D. Wittrup, Yeast surface display for screening combinatorial polypeptide libraries. Nat. Biotechnol. 15, 553–557 (1997).

38. S. Hochreiter, J. Schmidhuber, Long Short-Term Memory. Neural Comput. 9, 1735–1780 (1997).

39. D. M. Mason, S. Friedensohn, C. R. Weber, C. Jordi, B. Wagner, S. M. Meng, R. A. Ehling, L. Bonati, J. Dahinden, P. Gainza, B. E. Correia, S. T. Reddy, Optimization of therapeutic antibodies by predicting antigen specificity from antibody sequence via deep learning. Nat. Biomed. Eng., 1–13 (2021).

40. K. Saka, T. Kakuzaki, S. Metsugi, D. Kashiwagi, K. Yoshida, M. Wada, H. Tsunoda, R. Teramoto, Antibody design using LSTM based deep generative model from phage display library for affinity maturation. Sci. Rep. 11, 5852 (2021).

41. R. Akbar, P. A. Robert, C. R. Weber, M. Widrich, R. Frank, M. Pavlović, L. Scheffer, M. Chernigovskaya, I. Snapkov, A. Slabodkin, B. B. Mehta, E. Miho, F. Lund-Johansen, J. T. Andersen, S. Hochreiter, I. H. Haff, G. Klambauer, G. K. Sandve, V. Greiff, bioRxiv, in press, doi:10.1101/2021.07.08.451480.

42. E. Wilkinson, M. Giovanetti, H. Tegally, J. E. San, R. Lessells, D. Cuadros, D. P. Martin, D. A. Rasmussen, A.-R. N. Zekri, A. K. Sangare, A.-S. Ouedraogo, A. K. Sesay, A. Priscilla, A.-S. Kemi, A. M. Olubusuyi, A. O. O. Oluwapelumi, A. Hammami, A. A. Amuri, A. Sayed, A. E. O. Ouma, A. Elargoubi, N. A. Ajayi, A. F. Victoria, A. Kazeem, A. George, A. J. Trotter, A. A. Yahaya, A. K. Keita, A. Diallo, A. Kone, A. Souissi, A. Chtourou, A. V. Gutierrez, A. J. Page, A. Vinze, A. Iranzadeh, A. Lambisia, A. Ismail, A. Rosemary, A. Sylverken, A. Femi, A. Ibrahimi, B. Marycelin, B. S. Oderinde, B. Bolajoko, B. Dhaala, B. L. Herring, B.-M. Njanpop-Lafourcade, B. Kleinhans, B. McInnis, B. Tegomoh, C. Brook, C. B. Pratt, C. Scheepers, C. G. Akoua-Koffi, C. N. Agoti, C. Peyrefitte, C. Daubenberger, C. M. Morang’a, D. J. Nokes, D. G. Amoako, D. L. Bugembe, D. Park, D. Baker, D. Doolabh, D. Ssemwanga, D. Tshiabuila, D. Bassirou, D. S. Y. Amuzu, D. Goedhals, D. O. Omuoyo, D. Maruapula, E. Foster-Nyarko, E. K. Lusamaki, E. Simulundu, E. M. Ong’era, E. N. Ngabana, E. Shumba, E. El Fahime, E. Lokilo, E. Mukantwari, E. Philomena, E. Belarbi, E. Simon-Loriere, E. A. Anoh, F. Leendertz, F. Ajili, F. O. Enoch, F. Wasfi, F. Abdelmoula, F. S. Mosha, F. T. Takawira, F. Derrar, F. Bouzid, F. Onikepe, F. Adeola, F. M. Muyembe, F. Tanser, F. A. Dratibi, G. K. Mbunsu, G. Thilliez, G. L. Kay, G. Githinji, G. van Zyl, G. A. Awandare, G. Schubert, G. P. Maphalala, H. C. Ranaivoson, H. Lemriss, H. Anise, H. Abe, H. H. Karray, H. Nansumba, H. A. Elgahzaly, H. Gumbo, I. Smeti, I. B. Ayed, I. Odia, I. B. Ben Boubaker, I. Gaaloul, I. Gazy, I. Mudau, I. Ssewanyana, I. Konstantinus, J. B. Lekana-Douk, J.-C. C. Makangara, J.-J. M. Tamfum, J.-M. Heraud, J. G. Shaffer, J. Giandhari, J. Li, J. Yasuda, J. Q. Mends, J. Kiconco, J. M. Morobe, J. O. Gyapong, J. C. Okolie, J. T. Kayiwa, J. A. Edwards, J. Gyamfi, J. Farah, J. Nakaseegu, J. M. Ngoi, J. Namulondo, J. C. Andeko, J. J. Lutwama, J. O’Grady, K. Siddle, K. T. Adeyemi, K. A. Tumedi, K. M. Said, K. Hae-Young, K. O. Duedu, L. Belyamani, L. Fki-Berrajah, L. Singh, L. de O. Martins, L. Tyers, M. Ramuth, M. Mastouri, M. Aouni, M. el Hefnawi, M. I. Matsheka, M. Kebabonye, M. Diop, M. Turki, M. Paye, M. M. Nyaga, M. Mareka, M.-M. Damaris, M. W. Mburu, M. Mpina, M. Nwando, M. Owusu, M. R. Wiley, M. T. Youtchou, M. O. Ayekaba, M. Abouelhoda, M. G. Seadawy, M. K. Khalifa, M. Sekhele, M. Ouadghiri, M. M. Diagne, M. Mwenda, M. Allam, M. V. T. Phan, N. Abid, N. Touil, N. Rujeni, N. Kharrat, N. Ismael, N. Dia, N. Mabunda, N. Hsiao, N. B. Silochi, N. Nsenga, N. Gumede, N. Mulder, N. Ndodo, N. H. Razanajatovo, N. Iguosadolo, O. Judith, O. C. Kingsley, O. Sylvanus, O. Peter, O. Femi, O. Idowu, O. Testimony, O. E. Chukwuma, O. E. Ogah, C. K. Onwuamah, O. Cyril, O. Faye, O. Tomori, P. Ondoa, P. Combe, P. Semanda, P. E. Oluniyi, P. Arnaldo, P. K. Quashie, P. Dussart, P. A. Bester, P. K. Mbala, R. Ayivor-Djanie, R. Njouom, R. O. Phillips, R. Gorman, R. A. Kingsley, R. A. A. Carr, S. El Kabbaj, S. Gargouri, S. Masmoudi, S. Sankhe, S. B. Lawal, S. Kassim, S. Trabelsi, S. Metha, S. Kammoun, S. Lemriss, S. H. A. Agwa, S. Calvignac-Spencer, S. F. Schaffner, S. Doumbia, S. M. Mandanda, S. Aryeetey, S. S. Ahmed, S. Elhamoumi, S. Andriamandimby, S. Tope, S. Lekana-Douki, S. Prosolek, S. Ouangraoua, S. A. Mundeke, S. Rudder, S. Panji, S. Pillay, S. Engelbrecht, S. Nabadda, S. Behillil, S. L. Budiaki, S. van der Werf, T. Mashe, T. Aanniz, T. Mohale, T. Le-Viet, T. Schindler, U. J. Anyaneji, U. Chinedu, U. Ramphal, U. Jessica, U. George, V. Fonseca, V. Enouf, V. Gorova, W. H. Roshdy, W. K. Ampofo, W. Preiser, W. T. Choga, Y. Bediako, Y. Naidoo, Y. Butera, Z. R. de Laurent, A. A. Sall, A. Rebai, A. von Gottberg, B. Kouriba, C. Williamson, D. J. Bridges, I. Chikwe, J. N. Bhiman, M. Mine, M. Cotten, S. Moyo, S. Gaseitsiwe, N. Saasa, P. C. Sabeti, P. Kaleebu, Y. K. Tebeje, S. K. Tessema, C. Happi, J. Nkengasong, T. de Oliveira, A year of genomic surveillance reveals how the SARS-CoV-2 pandemic unfolded in Africa. Science. 374, 423–431 (2021).

43. C. Scheepers, J. Everatt, D. G. Amoako, A. Mnguni, A. Ismail, B. Mahlangu, C. K. Wibmer, E. Wilkinson, H. Tegally, J. E. San, J. Giandhari, N. Ntuli, S. Pillay, T. Mohale, Y. Naidoo, Z. Khumalo, Z. Makatini, N. for G. S. S. Africa (NGS-SA), A. Sigal, C. Williamson, F. Treurnicht, K. Mlisana, M. Venter, N. Hsiao, N. Wolter, N. Msomi, R. Lessells, T. Maponga, W. Preiser, P. L. Moore, A. von Gottberg, T. D. Oliveira, J. N. Bhiman, “The continuous evolution of SARS-CoV-2 in South Africa: a new lineage with rapid accumulation of mutations of concern and global detection” (2021), p. 2021.08.20.21262342,, doi:10.1101/2021.08.20.21262342.

44. J. Jumper, R. Evans, A. Pritzel, T. Green, M. Figurnov, O. Ronneberger, K. Tunyasuvunakool, R. Bates, A. Žídek, A. Potapenko, A. Bridgland, C. Meyer, S. A. A. Kohl, A. J. Ballard, A. Cowie, B. Romera-Paredes, S. Nikolov, R. Jain, J. Adler, T. Back, S. Petersen, D. Reiman, E. Clancy, M. Zielinski, M. Steinegger, M. Pacholska, T. Berghammer, S. Bodenstein, D. Silver, O. Vinyals, A. W. Senior, K. Kavukcuoglu, P. Kohli, D. Hassabis, Highly accurate protein structure prediction with AlphaFold. Nature. 596, 583–589 (2021).

45. B. M. J. W. van der Veer, J. Dingemans, L. B. van Alphen, C. J. P. A. Hoebe, P. H. M. Savelkoul, “A novel B.1.1.523 SARS-CoV-2 variant that combines many spike mutations linked to immune evasion with current variants of concern” (2021), p. 2021.09.16.460616,, doi:10.1101/2021.09.16.460616.

46. R. Antia, M. E. Halloran, Transition to endemicity: Understanding COVID-19. Immunity. 54, 2172–2176 (2021).

47. N. Phillips, The coronavirus is here to stay — here’s what that means. Nature. 590, 382–384 (2021).

48. D. Planas, D. Veyer, A. Baidaliuk, I. Staropoli, F. Guivel-Benhassine, M. M. Rajah, C. Planchais, F. Porrot, N. Robillard, J. Puech, M. Prot, F. Gallais, P. Gantner, A. Velay, J. Le Guen, N. Kassis-Chikhani, D. Edriss, L. Belec, A. Seve, L. Courtellemont, H. Péré, L. Hocqueloux, S. Fafi-Kremer, T. Prazuck, H. Mouquet, T. Bruel, E. Simon-Lorière, F. A. Rey, O. Schwartz, Reduced sensitivity of SARS-CoV-2 variant Delta to antibody neutralization. Nature. 596, 276–280 (2021).

49. T. N. Starr, N. Czudnochowski, Z. Liu, F. Zatta, Y.-J. Park, A. Addetia, D. Pinto, M. Beltramello, P. Hernandez, A. J. Greaney, R. Marzi, W. G. Glass, I. Zhang, A. S. Dingens, J. E. Bowen, M. A. Tortorici, A. C. Walls, J. A. Wojcechowskyj, A. De Marco, L. E. Rosen, J. Zhou, M. Montiel-Ruiz, H. Kaiser, J. R. Dillen, H. Tucker, J. Bassi, C. Silacci-Fregni, M. P. Housley, J. di Iulio, G. Lombardo, M. Agostini, N. Sprugasci, K. Culap, S. Jaconi, M. Meury, E. Dellota Jr, R. Abdelnabi, S.-Y. C. Foo, E. Cameroni, S. Stumpf, T. I. Croll, J. C. Nix, C. Havenar-Daughton, L. Piccoli, F. Benigni, J. Neyts, A. Telenti, F. A. Lempp, M. S. Pizzuto, J. D. Chodera, C. M. Hebner, H. W. Virgin, S. P. J. Whelan, D. Veesler, D. Corti, J. D. Bloom, G. Snell, SARS-CoV-2 RBD antibodies that maximize breadth and resistance to escape. Nature. 597, 97–102 (2021).

50. K. M. Hastie, H. Li, D. Bedinger, S. L. Schendel, S. M. Dennison, K. Li, V. Rayaprolu, X. Yu, C. Mann, M. Zandonatti, R. Diaz Avalos, D. Zyla, T. Buck, S. Hui, K. Shaffer, C. Hariharan, J. Yin, E. Olmedillas, A. Enriquez, D. Parekh, M. Abraha, E. Feeney, G. Q. Horn, CoVIC-DB team, Y. Aldon, H. Ali, S. Aracic, R. R. Cobb, R. S. Federman, J. M. Fernandez, J. Glanville, R. Green, G. Grigoryan, A. G. Lujan Hernandez, D. D. Ho, K.-Y. A. Huang, J. Ingraham, W. Jiang, P. Kellam, C. Kim, M. Kim, H. M. Kim, C. Kong, S. J. Krebs, F. Lan, G. Lang, S. Lee, C. L. Leung, J. Liu, Y. Lu, A. MacCamy, A. T. McGuire, A. L. Palser, T. H. Rabbitts, Z. Rikhtegaran Tehrani, M. M. Sajadi, R. W. Sanders, A. K. Sato, L. Schweizer, J. Seo, B. Shen, J. L. Snitselaar, L. Stamatatos, Y. Tan, M. T. Tomic, M. J. van Gils, S. Youssef, J. Yu, T. Z. Yuan, Q. Zhang, B. Peters, G. D. Tomaras, T. Germann, E. O. Saphire, Defining variant-resistant epitopes targeted by SARS-CoV-2 antibodies: A global consortium study. Science. 374, 472–478 (2021).

51. K. E. Kistler, T. Bedford, Evidence for adaptive evolution in the receptor-binding domain of seasonal coronaviruses OC43 and 229e. eLife. 10, e64509 (2021).

52. R. T. Eguia, K. H. D. Crawford, T. Stevens-Ayers, L. Kelnhofer-Millevolte, A. L. Greninger, J. A. Englund, M. J. Boeckh, J. D. Bloom, A human coronavirus evolves antigenically to escape antibody immunity. PLOS Pathog. 17, e1009453 (2021).

53. M. Worobey, J. Pekar, B. B. Larsen, M. I. Nelson, V. Hill, J. B. Joy, A. Rambaut, M. A. Suchard, J. O. Wertheim, P. Lemey, The emergence of SARS-CoV-2 in Europe and North America. Science. 370, 564–570 (2020).

## REFERENCES (METHODS)

1. T. N. Starr, A. J. Greaney, S. K. Hilton, D. Ellis, K. H. D. Crawford, A. S. Dingens, M. J. Navarro, J. E. Bowen, M. A. Tortorici, A. C. Walls, N. P. King, D. Veesler, J. D. Bloom, Deep Mutational Scanning of SARS-CoV-2 Receptor Binding Domain Reveals Constraints on Folding and ACE2 Binding. Cell. 182, 1295–1310.e20 (2020).

2. D. M. Mason, C. R. Weber, C. Parola, S. M. Meng, V. Greiff, W. J. Kelton, S. T. Reddy, High-throughput antibody engineering in mammalian cells by CRISPR/Cas9-mediated homology-directed mutagenesis. Nucleic Acids Res. 46, 7436–7449 (2018).

3. E. T. Boder, K. D. Wittrup, Yeast surface display for screening combinatorial polypeptide libraries. Nat. Biotechnol. 15, 553–557 (1997).

4. G. Chao, W. L. Lau, B. J. Hackel, S. L. Sazinsky, S. M. Lippow, K. D. Wittrup, Isolating and engineering human antibodies using yeast surface display. Nat. Protoc. 1, 755–768 (2006).

5. R. Vazquez-Lombardi, D. Nevoltris, A. Luthra, P. Schofield, C. Zimmermann, D. Christ, Transient expression of human antibodies in mammalian cells. Nat. Protoc. 13, 99–117 (2018).

6. R Core Team, R: A Language and Environment for Statistical Computing (R Foundation for Statistical Computing, Vienna, Austria; https://www.R-project.org).

7. G. V. Rossum, F. L. Jr. Drake, The Python Language Reference Manual (Network Theory Ltd, 2011).

8. H. Wickham, ggplot2: Elegant Graphics for Data Analysis (Springer-Verlag New York, 2009; http://ggplot2.org).

9. Z. Gu, R. Eils, M. Schlesner, Complex heatmaps reveal patterns and correlations in multidimensional genomic data. Bioinforma. Oxf. Engl. 32, 2847–2849 (2016).

10. R. Kolde, pheatmap: Pretty Heatmaps (2019; https://CRAN.R-project.org/package=pheatmap).

11. G. Csardi, T. Nepusz, The igraph software package for complex network research. InterJournal. Complex Systems, 1695 (2006).

12. J. A. Gustavsen, S. Pai, R. Isserlin, B. Demchak, A. R. Pico, RCy3: Network biology using Cytoscape from within R (2019),, doi:10.12688/f1000research.20887.3.

13. H. Wickham, stringr: Simple, Consistent Wrappers for Common String Operations (2019; https://CRAN.R-project.org/package=stringr).

14. H. Wickham, R. François, L. Henry, K. Müller, dplyr: A Grammar of Data Manipulation (2021; https://CRAN.R-project.org/package=dplyr).

15. E. Neuwirth, RColorBrewer: ColorBrewer Palettes (2014; https://CRAN.R-project.org/package=RColorBrewer).

16. P. Shannon, A. Markiel, O. Ozier, N. S. Baliga, J. T. Wang, D. Ramage, N. Amin, B. Schwikowski, T. Ideker, Cytoscape: a software environment for integrated models of biomolecular interaction networks. Genome Res. 13, 2498–2504 (2003).

17. J. Platt, Probabilistic Outputs for Support Vector Machines and Comparisons to Regularized Likelihood Methods. Adv Large Margin Classif. 10 (2000).

18. H. Boström, in 2008 Seventh International Conference on Machine Learning and Applications (IEEE, San Diego, CA, USA, 2008; http://ieeexplore.ieee.org/document/4724964/), pp. 121–126.

19. A. Niculescu-Mizil, R. Caruana, in Proceedings of the 22nd international conference on Machine learning - ICML ‘05 (ACM Press, Bonn, Germany, 2005; http://portal.acm.org/citation.cfm?doid=1102351.1102430), pp. 625–632.

20. J. Jumper, R. Evans, A. Pritzel, T. Green, M. Figurnov, O. Ronneberger, K. Tunyasuvunakool, R. Bates, A. Žídek, A. Potapenko, A. Bridgland, C. Meyer, S. A. A. Kohl, A. J. Ballard, A. Cowie, B. Romera-Paredes, S. Nikolov, R. Jain, J. Adler, T. Back, S. Petersen, D. Reiman, E. Clancy, M. Zielinski, M. Steinegger, M. Pacholska, T. Berghammer, S. Bodenstein, D. Silver, O. Vinyals, A. W. Senior, K. Kavukcuoglu, P. Kohli, D. Hassabis, Highly accurate protein structure prediction with AlphaFold. Nature. 596, 583–589 (2021).

21. L. Schrödinger, W. DeLano, PyMOL (2020; http://www.pymol.org/pymol).

